# QuICSeedR: An R package for analyzing fluorophore-assisted seed amplification assay data

**DOI:** 10.1101/2024.08.27.609919

**Authors:** Manci Li, Damani N. Bryant, Sarah Gresh, Marissa Milstein, Peter R. Christenson, Stuart S. Lichtenberg, Peter A. Larsen, Sang-Hyun Oh

**Author notes:** **Corresponding author** Manci Li, PhD.

## Abstract

**Summary:** Fluorophore-assisted seed amplification assays (F-SAAs), such as real-time quaking-induced conversion (RT-QuIC) and fluorophore-assisted protein misfolding cyclic amplification (F-PMCA), have become indispensable tools for studying protein misfolding in neurodegenerative diseases. However, analyzing data generated by these techniques often requires complex and time-consuming manual processes. Additionally, the lack of standardization in F-SAA data analysis presents a significant challenge to the interpretation and reproducibility of F-SAA results across different laboratories and studies. Here, we present QuICSeedR (pronounced as “quick seeder”), an R package that addresses these challenges by providing a comprehensive toolkit for the automated processing, analysis, and visualization of F-SAA data. Importantly, QuICSeedR also sets up the foundation for building an F-SAA data management and analysis framework, enabling more consistent and comparable results across different research groups.

**Availability and implementation:** QuICSeedR source code is freely available at: https://github.com/mancili/QuICSeedR. Data and code used in this manuscript are provided in Supplementary Materials.

**Supplementary information:** Supplementary Materials are available with the manuscript.

## 1 Introduction

Neurodegenerative diseases (NDDs), such as Alzheimer’s disease, Parkinson’s disease (PD), multiple system atrophy (MSA), amyotrophic lateral sclerosis, and prion diseases (e.g. Creutzfeldt-Jakob disease in humans and chronic wasting disease in cervids), are characterized by protein misfolding and aggregation processes that pose significant challenges for early diagnosis, accurate prognosis, and effective treatment (Duncan, 2011; Hou *et al*., 2019; Dokholyan *et al*., 2022). Pathological protein misfolding and accumulation (i.e., amyloid formation) start decades before clinical signs (Prusiner, 2001; Soto and Pritzkow, 2018).

Fluorophore-assisted seed amplification assays (F-SAAs), such as real-time quaking-induced conversion (RT-QuIC) (Wilham *et al*., 2010; Atarashi *et al*., 2011; Zanusso *et al*., 2016; Metrick *et al*., 2019; Candelise *et al*., 2020) and fluorophore-assisted protein misfolding cyclic amplification (F-PMCA) (Shahnawaz *et al*., 2017, 2020; Singer *et al*., 2020, 2021), have emerged as powerful tools for detecting misfolded protein aggregates with high sensitivity and specificity (Salvadores *et al*., 2014; Scialò *et al*., 2020; Coysh and Mead, 2022; Brockmann *et al*., 2024; Shahnawaz *et al*., 2020). Recent advancements in fluorophore-assisted QuIC techniques, such as Micro-QuIC and Nano-QuIC, have further improved reaction kinetics, enabling faster detection and potentially higher sensitivity in clinical applications (Lee *et al*., 2024; Christenson *et al*., 2023). These assays leverage the ability of misfolded proteins to induce conformational changes in normally folded proteins while providing an optimized environment that accelerates this process, which can be monitored in real-time using a fluorophore (e.g. thioflavin T, ThT) structurally sensitive to amyloid (Vassar and Culling, 1959; Naiki *et al*., 1989; Biancalana and Koide, 2010; Gade Malmos *et al*., 2017). The resulting ultrasensitivity holds significant promise for large-scale clinical applications, revolutionizing early diagnosis and monitoring of disease progression for NDDs (Coysh and Mead, 2022).

Despite their utility, the analysis of data from these assays presents several challenges. First, experiments often involve multiple 96- or 384-well plates with time-step fluorescence readings over days, generating substantial amounts of data that are time-consuming to process manually. This necessitates a computable and automation-friendly data framework to streamline analysis. Second, assays may differ in plate layout, sample types, and analysis requirements, needing flexible analytical tools that can adapt to various experimental designs. In addition, different analysis methods may be needed for various sample types or experimental conditions, requiring workflow that facilitates method comparison. These challenges underscore the need for domain-specific data management and analysis solutions that can handle high-volume, diverse, and complex experimental data efficiently and accurately.

Here, we developed QuICSeedR, an R-based toolkit, to address these challenges by providing recommendations for efficient data management framework, automating data processing, supporting large-scale analysis, and enabling comparative studies of analysis methods.

## 2 Implementation

The QuICSeedR workflow processes two primary inputs (Fig. 1): 1) plate data containing experimental setup and sample placement information and 2) time-course fluorescence data, typically exported from MARS, the Microplate Reader Software used by BMG LABTECH plate readers (BMG LABTECH) (the current standard for F-SAAs). The package can, however, accommodate any time-series data. The cleaning phase integrates and processes these inputs. The calculation stage computes commonly used metrics in F-SAA research, such as rate of amyloid formation (RAF) (Henderson *et al*., 2015), maximum slope (MS) (Haley *et al*., 2013; Gallups and Harms, 2022), and max-point ratio (MPR) (Rowden *et al*., 2023), with options for normalization (Li *et al*., 2021; Christenson *et al*., 2023). Results are then reformatted to ensure compatibility with visualization tools in popular graphing software, such as GraphPad Prism (GraphPad) and OriginLab (OriginLab). The analysis phase conducts statistical tests and inter-group comparisons, followed by a result summarization phase (Fig. 1). QuICSeedR also offers diverse visualization options in addition to GraphPad Prism integration throughout, streamlining the processing and analysis of complex assay data to achieve robust and reproducible research in protein misfolding and aggregation studies (Fig. 1). In addition, QuICSeedR provides batch processing capabilities to support large-scale analysis (Fig. 1). In the following sections, we present implementations of QuICSeedR, demonstrating its utility across different sample types, F-SAAs, and NDDs in both humans and animals.

**Fig. 1.**
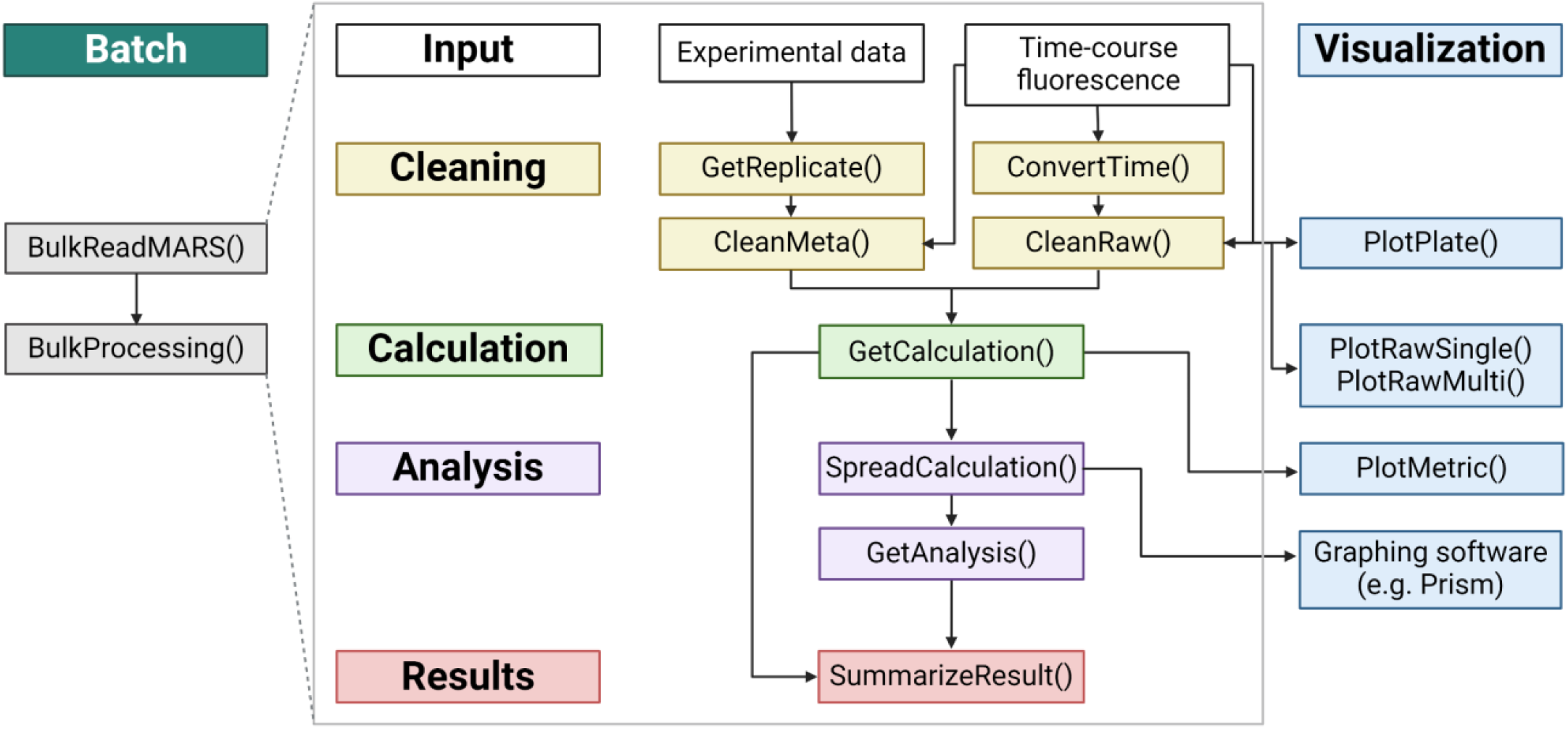
Workflow of QuICSeedR for fluorophore-assisted seed amplification assays (F-SAA). This diagram outlines the streamlined process for analyzing F-SAA data. The workflow progresses through key stages: input of experimental data, cleaning (data preprocessing), calculation, analysis, results generation, and visualization. It includes functions for batch processing, time-course data handling, and various data cleaning, calculation, and plotting operations. This workflow is designed to enhance efficiency across diverse experimental designs in F-SAA studies by providing a cohesive framework for data management, analysis, and visualization.

### 2.1 Improved efficiency by unified data architecture for diverse experimental designs

QuICSeedR introduces a unified data architecture that significantly boosts efficiency across diverse experimental designs in F-SAAs. This architecture is designed to make experimental data of F-SAA computable; it utilizes plate layout files for experimental setup documentation and files for raw fluorescence data (Supplemental Material). The latter is exported from the MARS software used by BMG LABTECH plate readers (the current standard instrument for F-SAAs) (BMG LABTECH; Atarashi *et al*., 2011; Shahnawaz *et al*., 2017). While optimized for MARS output, QuICSeedR demonstrates flexibility in accommodating data from alternative instruments. Users can easily adapt data from other sources to fit the required format, ensuring broad applicability across different experimental setups and instrumentation.

At the reaction level, the package offers granular control over data processing. Users can split sample identifiers into multiple variables as needed, with the flexibility to add additional variables after initial processing (Supplemental Material). This adaptability ensures that the data structure can accommodate a wide range of experimental designs and evolving research needs. Further, the unified architecture seamlessly handles both 96- and 384-well plate formats, providing a versatile platform for multiple experimental scales.

Overall, using QuICSeedR and implementing unified data architecture resulted in a significant reduction in analysis time, with processing duration decreasing from 1-2 hours of manual effort per plate to less than one minute of automated processing (Supplementary Materials).

### 2.2 Enhanced analytical power through the integration of literature-based statistical and visualization options

QuICSeedR incorporates a suite of statistical analyses and visualization tools derived from established literature in F-SAA and protein misfolding research. This integration provides researchers with convenient access to commonly used analytical methods, such as threshold determination (e.g., standard deviation-based, background ratio-based, or relative fluorescence unit cut-off) (Wilham *et al*., 2010; Cheng *et al*., 2016; Shahnawaz *et al*., 2020), kinetic parameter calculations (e.g., time to threshold, rate of amyloid formation, maximum slope, and maximum point ratio) (Henderson *et al*., 2015; Li *et al*., 2021; Haley *et al*., 2013; Gallups and Harms, 2022; Rowden *et al*., 2023), and statistical analyses between sample groups.

The visualization options are designed to align with existing research. They include time-course fluorescence plots for plate layouts (Supplementary Materials), multiple samples colored by sample (Fig. 2A), and single samples colored by technical replicates (Fig. 2B), as well as key metrics across different experimental conditions compatible with customization using the ggplot2 package (Fig. 2C) (Wickham, 2016).

**Fig. 2.**
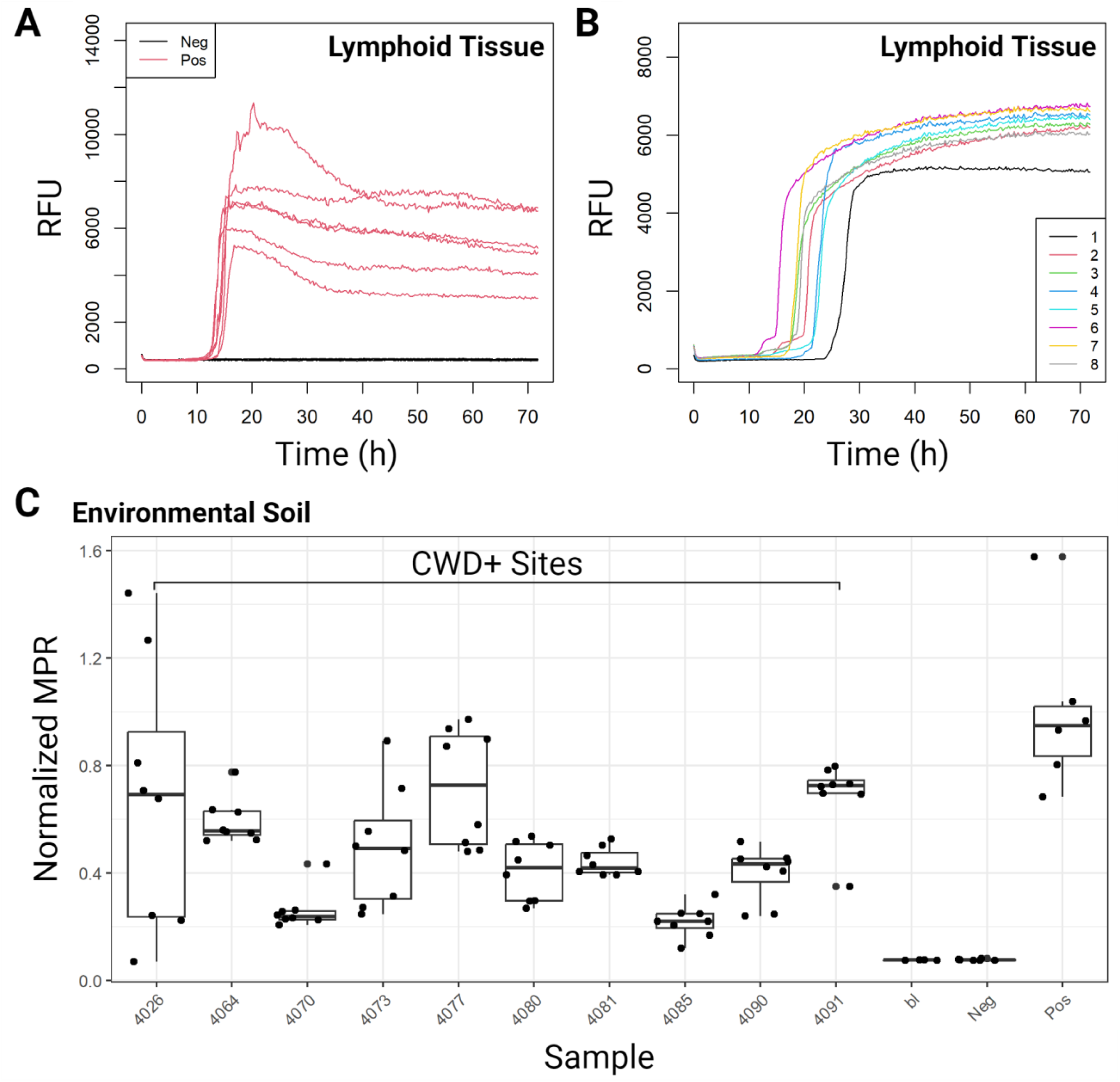
Exemplary visualization options in QuICSeedR. **(A)** Time-course fluorescence (measurements taken every 15 minutes) of multiple samples from a 96-well experiment, comparing positive and negative lymphoid tissues. **(B)** Single-sample time-course fluorescence from a 384-well plate, showing replicates of positive lymphoid tissue. **(C)** Max-point ratio (MPR) metric plot of environmental soil samples from a chronic wasting disease-positive site.

### 2.3 Achieving scalability by high-throughput analysis

QuICSeedR leverages list structures in R to facilitate efficient data storage and retrieval, enabling high-throughput analysis (R Core Team). We demonstrated QuICSeedR’s high-throughput capability to handle complex, multi-dimensional datasets typical of extensive F-SAA experiments by applying it to a subset of data from a large-scale chronic wasting disease (CWD) environmental swab study (Milstein *et al*., 2024). In this application, QuICSeedR’s automated workflow read, processed, and analyzed data from 242 samples, encompassing 1,152 individual reactions, each comprising 65-95 time-steps of fluorescence readings (Supplementary Materials). Remarkably, this automated process completed in seconds what would have required 8-24 hours of manual labor, representing a reduction of over 99% in processing time. A subset of the data was visualized in Fig. 3 (Milstein *et al*., 2024).

**Fig. 3.**
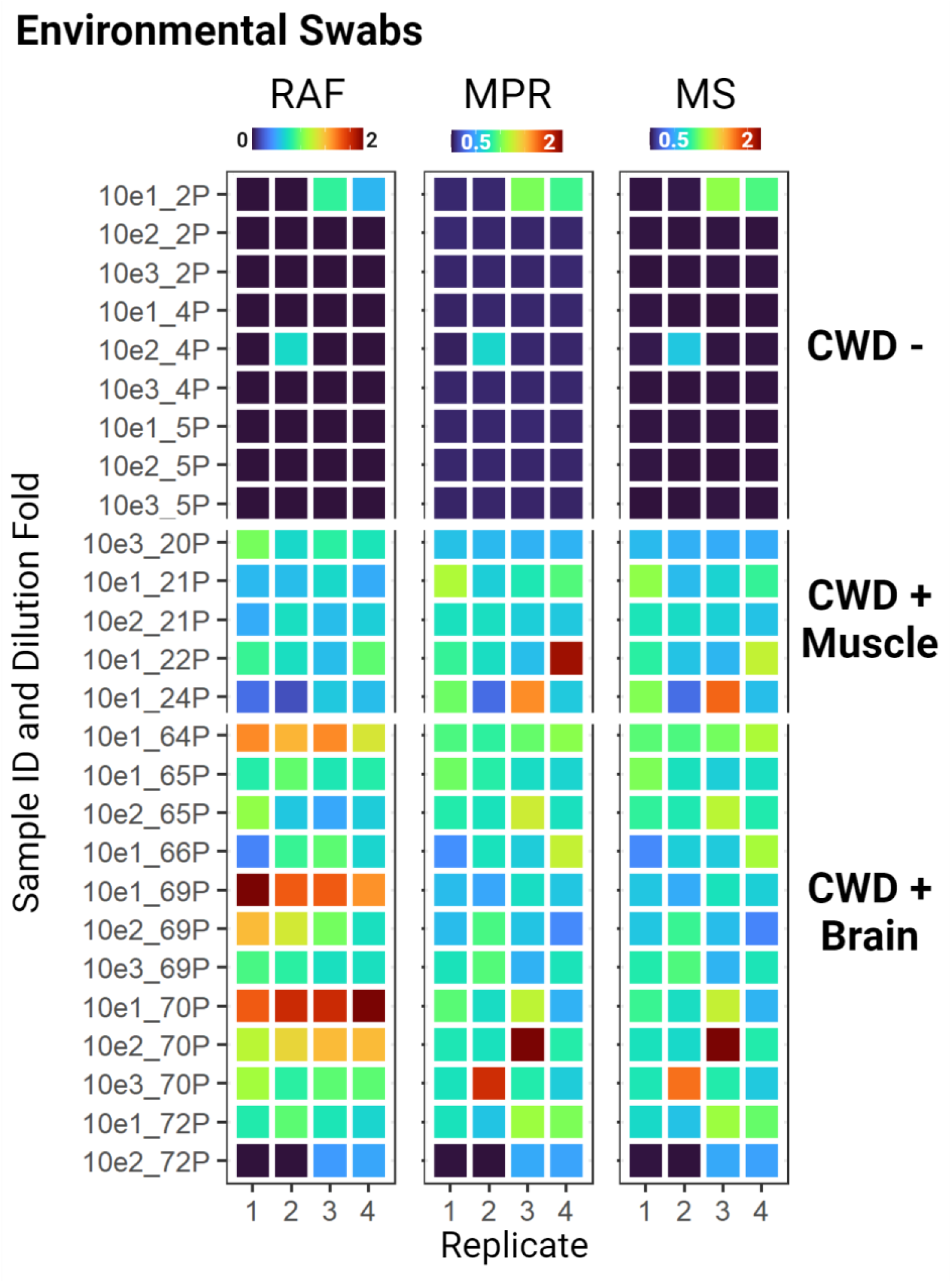
High-throughput capability of QuICSeedR. Normalized metrics (RAF, MPR, MS) from a subset of the 242 samples and 1,152 RT-QuIC reactions are visualized. These samples are equipment swabs from various tissue processing scenarios. Samples 10e1_2P through 10e3_5P represent swabs from equipment that processed CWD-negative muscle tissue. Samples 10e3_20P through 10e1_24P demonstrated significant seeding activity in RT-QuIC assays and were of different dilutions of swabs from equipment that processed CWD-positive muscles. Samples 10e1_64P to 10e2_72P showed significant seeding activity in RT-QuIC assays and were different dilutions of swabs from equipment used to process CWD-positive brain samples. RAF, rate of amyloid formation; MPR, max-point ratio; MS, maximum slope; RT-QuIC, real-time quaking-induced conversion; 10e1, 10-fold dilution; 10e2, 100-fold dilution; 10e3, 1000-fold dilution.

### 2.4 Versatile applications to different NDDs, sample types, and F-SAAs

The versatility of QuICSeedR extends beyond its data-handling capabilities to its applicability across various NDDs, sample types, and assays. In addition to analyzing CWD prion seeding activity in RT-QuIC in both 96- and 384-well formats for environmental soil samples (Fig. 2; Supplementary Materials), we applied QuICSeedR to data derived from other diverse arrays of sample matrices, including environmental surface swabs (Fig. 3) as well as various biological tissues. Specifically, the package has shown utility in analyzing samples relevant to CWD diagnosis, monitoring, and surveillance, including ear punch tissue from elk (*Cervus canadensis*), as well as lymphoid, blood, and muscle tissues from white-tailed deer (*Odocoileus virginianus*) (Fig. 4A; Supplementary Materials) (Schwabenlander *et al*., 2022; Li *et al*., 2021; Bryant *et al*., 2024).

**Fig. 4.**
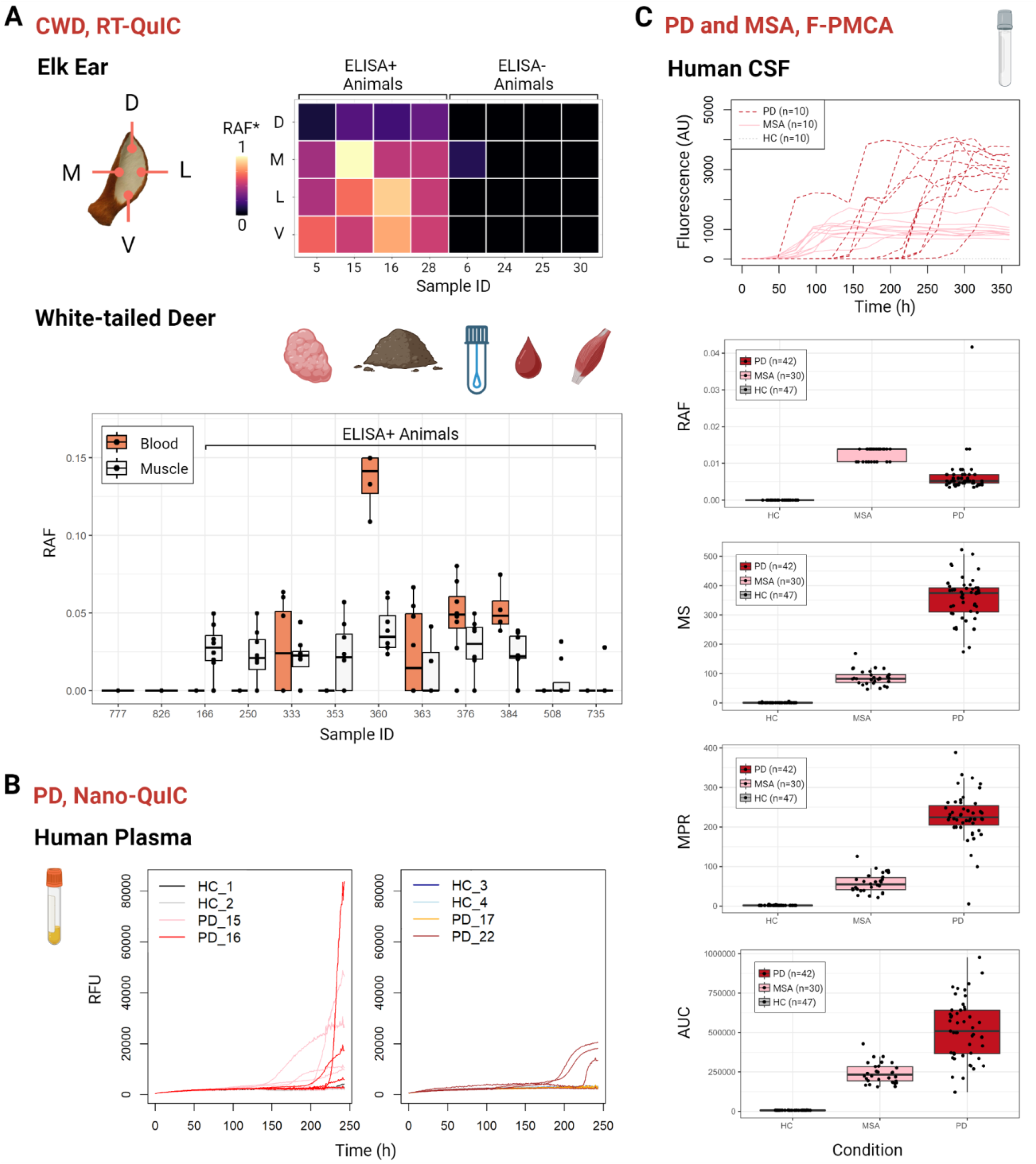
Versatility of QuICSeedR across neurodegenerative diseases, species, sample types, and fluorophore-assisted seed amplification assays. **(A)** Chronic wasting disease (CWD) in RT-QuIC. Normalized rate of amyloid formation (RAF) values from different ear sections of ELISA+ and ELISA-elk were shown. QuICSeedR achieved analysis and visualization for RT-QuIC data generated from lymphoid tissues, soil, swabs, blood, and muscle samples in white-tailed deer. RAF values in blood and muscle samples from ELISA+ white-tailed deer are highlighted in a boxplot. RT-QuIC, real-time quaking-induced conversion; RAF*, normalized RAF; ELISA, enzyme-linked immunosorbent assay; D, dorsal; V, ventral; L, lateral; M, medial. **(B)** Detecting misfolded ɑ-synuclein seeding activity in human plasma from PD using nano-QuIC. HC, healthy control; PD, Parkinson’s diseases; nano-QuIC, nanoparticle-enhanced RT-QuIC. **(C)** Detecting misfolded ɑ-synuclein seeding activity in human cerebrospinal fluid (CSF) from PD and multiple system atrophy (MSA) patients using fluorophore-assisted protein misfolding cyclic amplification assay (F-PMCA). Wilcoxon rank-sum tests were performed to compare the relative amplification factor (RAF), max-point ratio (MPR), maximum slope, and area under the curve (AUC) between each pair of groups: Parkinson’s disease (PD) patients and control individuals, multiple system atrophy (MSA) patients and control individuals, and PD patients and MSA patients. All comparisons resulted in p-values less than 0.0001.

Furthermore, QuICSeedR can be used to analyze data from different assay types and NDDs. We showcased its application using two curated datasets: plasma nanoparticle-enhanced RT-QuIC (nano-QuIC) data from PD patients and controls (Christenson *et al*., 2023) (Fig. 4B; Supplementary Materials), alongside cerebrospinal fluid F-PMCA data from PD and MSA patients, as well as controls (Shahnawaz *et al*., 2020) (Fig. 4C; Supplementary Materials). Both assays aimed to detect α-synuclein seeding activity in these synucleinopathies (Supplementary Materials). This cross-assay compatibility underscores the package’s potential to integrate and compare results from various protein misfolding detection techniques.

## Discussion

QuICSeedR represents a significant advancement in F-SAA data management and analysis, addressing key limitations in existing software: it streamlines workflows and facilitates comprehensive experimental designs in protein misfolding research by automating data processing, supporting large-scale analysis, and enabling method comparisons. The unified approach of QuICSeedR to diverse applications—from various plate formats to different sample types and assay methods—improves efficiency in individual experiments and cross-study comparisons. Its consolidation of analytical and visualization capabilities promotes standardization and reproducibility of F-SAAs across laboratories, allowing easier result comparisons and more consistent reporting. Such comparative analysis and standardization are needed for effective translation of F-SAA for both animal and human NDDs (Pétavy *et al*., 2019; Barros *et al*., 2022).

While QuICSeedR is currently optimized for data output from BMG LABTECH instruments, its modular design allows for potential expansion. The toolkit can be adapted to incorporate functions that read data from various hardware platforms and different assay types outside of F-SAAs. Such extensions could encompass ThT fluorescence assays, HANABI, and other fluorescence and colorimetric implementations (Naiki *et al*., 1989; Ogi *et al*., 2014; Goto *et al*., 2022). Additionally, QuICSeedR’s flexible structure can accommodate data from other assays important for detecting or studying misfolded proteins, such as ELISA (NVAP reference guide: Chronic wasting disease (control and eradication)). This inherent adaptability positions QuICSeedR to evolve alongside the changing needs of researchers across multiple experimental paradigms, should future development be pursued.

QuICSeedR’s structured workflow supports organized data handling, laying the groundwork for a comprehensive data repository that could enhance reproducibility and collaborative research efforts in F-SAA studies. This forward-thinking approach will not only facilitate data sharing and reuse but also enable seamless integration with machine learning applications, allowing for advanced pattern recognition, predictive modeling, and automated feature extraction from F-SAA data in the future. Ultimately, QuICSeedR is positioned to accelerate the translation of F-SAA into high-throughput clinical applications—such as early diagnosis protocols, patient selection for clinical trials, and monitoring treatment efficacy—for both animal and human NDDs.

## Conclusion

QuICSeedR addresses the current analytical needs of F-SAA and anticipates future data management requirements in the field. As the field of protein misfolding research continues to evolve, tools like QuICSeedR will play an instrumental role in accelerating scientific discoveries and their translation into clinical practice.

## Author Contributions

ML conceived and designed the study, developed the software, performed data analysis and visualization, and drafted the manuscript. DB, SG, MM, and PC contributed to data curation. SHO and PAL reviewed and edited the manuscript. All authors reviewed and approved the final version of the manuscript.

## Acknowledgments

We extend our sincere gratitude to Dr. Catalina Picasso for her valuable preliminary review of QuICSeedR, and to Madeline Grunklee and Corina Valencia Tibbitts for their helpful feedback during initial testing. We also acknowledge with appreciation the insights provided by Dr. Jeremy Chacon from the Minnesota Supercomputing Institute regarding the CRAN submission process, and the advice on biostatistics offered by Dr. Aaron Rendahl. We thank Dr. Wolfgang Singer, Dr. Pinaki Misra, and Shannen Griffiths for their helpful comments. The figures in this manuscript were created with the assistance of BioRender.

