## Supplementary material for "QuICSeedR: An R package for analyzing fluorophore-assisted seed amplification assay data": QuICSeedR_0.1.2.pdf

### Package ‘QuICSeedR’

August 27, 2024

**Title** Analyze Data for Fluorophore-Assisted Seed Amplification Assays

**Date** 2024-08-27

**Version** 0.1.2

**Depends** R (>= 4.1.0)

**Copyright** Manci Li

**Description** A toolkit for analysis and visualization of data from fluorophore-assisted seed amplification assays, such as Real-Time Quaking-Induced Conversion (RT-QuIC) and Fluorophore-Assisted Protein Misfolding Cyclic Amplification (PMCA). 'QuICSeedR' addresses limitations in existing software by automating data processing, supporting large-scale analysis, and enabling comparative studies of analysis methods. It incorporates methods described in Henderson et al. (2015) <doi:10.1099/vir.0.069906-0>, Li et al. (2020) <doi:10.1038/s41598-021-96127-8>, Rowden et al. (2023) <doi:10.3390/pathogens12020309>, Haley et al. (2013) <doi:10.1371/journal.pone.0081488>, and Mair and Wilcox (2020) <doi:10.3758/s13428-019-01246-w>. Please refer to the original publications for details.

**Imports** WRS2, magrittr, graphics, stats, ggplot2, methods, dplyr, tidyr, tidyselect, rlang, readxl

**License** GPL-3 | file LICENSE

**Encoding** UTF-8

**Roxygen** list(markdown = TRUE)

**RoxygenNote** 7.3.2

**Suggests** testthat (>= 3.0.0)

**Config/testthat/edition** 3

**Index** **20**

---

`BulkProcessing` *Processing and analyzing Multiple Experiments*

---

#### Description

This function processes multiple experiments in parallel, performing a series of operations including time conversion, metadata cleaning, raw data cleaning, calculations, analysis, and result summarization.

#### Usage

```
BulkProcessing(data, do_analysis = TRUE, params = list(), verbose = FALSE)
```

#### Arguments

| | |
|---|---|
| `data` | Compiled data of experiments. |
| `do_analysis` | Logical. Whether statistic analysis is included. Default is `TRUE`. |
| `params` | A list of parameters for various processing steps. If a parameter is not provided, default values will be used. `control` used in `GetAnalysis()` is required. parameters can be provided for the following functions: <br>• `ConvertTime`<br>• `CleanMeta`<br>• `CleanRaw`<br>• `GetCalculation`<br>• `SpreadCalculation`<br>• `GetAnalysis`<br>• `SummarizeResult` |
| `verbose` | Logical; if TRUE, prints detailed processing information for troubleshooting. Default is FALSE. |

#### Value

A list containing three elements:

- combined_calculation: A data frame of combined calculations from all experiments
- combined_cleanraw: Cleaned raw data for each experiment
- combined_result: A data frame of combined results from all experiments

#### Examples

```
#Get path for example data
path = system.file("extdata", package = "QuICSeedR")

#Helper function
add_underscore <- function(text) {
  gsub("([a-zA-Z])(\\d)", "\\1_\\2", text)
  }

#Read in data
elkear = BulkReadMARS(path = path,
                      plate_subfix = 'plate',
                      raw_subfix = 'raw',
                      helper_func = add_underscore)

#Set up parameters for batch analysis
params = list(
  CleanMeta = list(split_content = TRUE, split_into = c('region', 'sample')),
  GetCalculation = list(cycle_background = 5, norm = TRUE, norm_ct = 'Pos',
                       sd_fold = 10, time_skip = 5),
  GetAnalysis = list(control = "Neg")
  )

#Get results
results = BulkProcessing(data =elkear, params = params)

str(results)
```

---

BulkReadMARS *Bulk Read MARS data*

---

#### Description

Bulk Read MARS data

#### Usage

```
BulkReadMARS(path, plate_subfix, raw_subfix, helper_func = NULL)
```

#### Arguments

| | |
|---|---|
| path | Character string specifying the path to the parent directory containing the data folders. |
| plate_subfix | Character string used to identify plate data files. |
| raw_subfix | Character string used to identify raw data files. |
| helper_func | Optional function to be applied to each column of the plate data (default: NULL). |

##### Details

The example dataset, located in `inst/extdata/`, consists of folders representing individual experimental runs. This dataset is a subset of the data used in a publication led by Dr. Stuart Lichtenberg. For detailed information about the experiments, please contact Dr. Stuart Lichtenberg at ``.

Each folder contains two Excel files:

- A file with the suffix `plate`: Contains the plate setup and experimental information.
- A file with the suffix `raw`: Contains fluorescence data exported from MARS software.

##### Value

A list containing data from each folder, including plate, raw, and replicate data.

##### Examples

```
path = system.file("extdata", package = "QuICSeedR")
data <- BulkReadMARS(path = path,
                     plate_subfix = 'plate',
                     raw_subfix = 'raw')
str(data)
```

---

#### CleanMeta *Get Clean Metadata*

---

##### Description

This function processes raw data (raw), plate layout (plate), and replicate information (replicate) to create a clean metadata dataframe. It can optionally split the content column into additional columns.

##### Usage

```
CleanMeta(
  raw,
  plate,
  replicate,
  split_content = FALSE,
  split_by = "_",
  split_into = c(),
  del_na = TRUE
)
```

##### Arguments

| | |
|---|---|
| `raw` | A dataframe containing the raw data. |
| `plate` | A dataframe containing the plate layout information. |
| `replicate` | A dataframe containing the replicate information. Output of `GetReplicate()`. |
| `split_content` | Logical, whether to split the content. Default is FALSE. |

| | |
|---|---|
| split_by | A character string to split the content by. Default is "_". |
| split_into | A character vector specifying names for the split columns. Required if split_content is TRUE. |
| del_na | Logical, whether to drop rows containing NA. Default is TRUE. |

##### Value

A data frame containing the cleaned metadata.

##### Examples

```
# Define the path to the plate data file
plate_path <- system.file("extdata/20240716_p3",
                          file = '20240716_p3_plate.xlsx',
                          package = "QuICSeedR")

# Read the plate data
plate <- readxl::read_xlsx(plate_path)

# Define the path to the raw data file
raw_path <- system.file("extdata/20240716_p3",
                        file = '20240716_p3_raw.xlsx',
                        package = "QuICSeedR")
# Read the raw data
raw <- readxl::read_xlsx(raw_path)

# Get replicate data
replicate <- GetReplicate(plate)

#Get metadata and display the few rows
meta = CleanMeta(raw, plate, replicate)
head(meta)
```

---

#### CleanRaw *Generate Clean Raw Data*

---

##### Description

This function takes metadata, raw data, and total cycle information to generate clean raw fluorescence data.

##### Usage

```
CleanRaw(meta, raw, plate_time, cycle_total)
```

##### Arguments

| | |
|---|---|
| meta | A data frame containing the metadata. Output from CleanMeta function. |
| raw | Raw fluorescence readings from MARS software. |
| plate_time | Output of ConvertTime(). |
| cycle_total | The total number of cycles (rows) to include in the output. |

##### Value

A data frame containing the cleaned raw fluorescence data.

##### Examples

```
# Define the path to the plate data file
plate_path <- system.file("extdata/20240716_p3",
                          file = '20240716_p3_plate.xlsx',
                          package = "QuICSeedR")

# Read the plate data
plate <- readxl::read_xlsx(plate_path)

# Define the path to the raw data file
raw_path <- system.file("extdata/20240716_p3",
                        file = '20240716_p3_raw.xlsx',
                        package = "QuICSeedR")
# Read the raw data
raw <- readxl::read_xlsx(raw_path)

# Get replicate data
replicate <- GetReplicate(plate)

# Ensure time displayed as decimal hours
plate_time = ConvertTime(raw)

#Get metadata and display the few rows
meta = CleanMeta(raw, plate, replicate)

#Clean data
cleaned_data <- CleanRaw(meta, raw, plate_time)

cleaned_data[1:5, 1:5]
```

---

#### `ConvertTime` *Extract and Convert Time Data to Decimal Hours*

---

##### Description

The function extracts and converts run time information from MARS output.

##### Usage

```
ConvertTime(raw)
```

##### Arguments

| | |
|---|---|
| `raw` | Output from MARS. |

##### Value

A data frame containing the time information in decimal hours.

#### Examples

```
# Example with "hours and minutes" format
raw_data <- data.frame(
  V1 = c("Header", "Row1", "Row2"),
  V2 = c("Time", "1 h 30 min", "2 h 45 min")
)
plate_time <- ConvertTime(raw_data)
print(plate_time)

# Example with decimal hours format
raw_data2 <- data.frame(
  V1 = c("Header", "Row1", "Row2"),
  V2 = c("Time", "1.5", "2.75")
)
plate_time2 <- ConvertTime(raw_data2)
print(plate_time2)
```

---

GetAnalysis *Perform Statistical Analysis on Calculations*

---

#### Description

This function performs statistical analysis on a list of calculation spreads, comparing each column to a control column using various statistical tests.

#### Usage

```
GetAnalysis(
  calculation_spread,
  control,
  test = "wilcox",
  alternative = "two.sided",
  adjust_p = FALSE,
  alpha = 0.05
)
```

#### Arguments

`calculation_spread`
: A list of data frames, each representing a calculation spread.
  Output of `SpreadCaculation()`.

`control`
: The name or pattern of the control column in each data frame.

`test`
: The statistical test to use. Options are:
  - "t-test": Student's t-test, suitable for normally distributed data. For more information, run: `?stats::t.test`
  - "wilcox": Wilcoxon rank-sum test (also known as Mann-Whitney U test), a non-parametric test. For more information, run: `?stats::wilcox.test`
  - "yuen": Yuen's test for trimmed means, robust against outliers and non-normality. For more information, run: `?WRS2::yuen`

Default is "wilcox".

alternative — Options are "two.sided", "less", or "greater". Default is "two.sided".

adjust_p — Logical. Whether to adjust p-values for multiple comparisons. Default is FALSE.

alpha — The significance level for determining significance stars. Default is 0.05.

##### Value

A list of data frames containing the results of the statistical analysis for selected metrics.

##### Examples

```
# Define the path to the plate data file
plate_path <- system.file("extdata/20240716_p3",
                          file = '20240716_p3_plate.xlsx',
                          package = "QuICSeedR")

# Read the plate data
plate <- readxl::read_xlsx(plate_path)

# Define the path to the raw data file
raw_path <- system.file("extdata/20240716_p3",
                        file = '20240716_p3_raw.xlsx',
                        package = "QuICSeedR")
# Read the raw data
raw <- readxl::read_xlsx(raw_path)

# Get replicate data
replicate <- GetReplicate(plate)

# Ensure time displayed as decimal hours
plate_time = ConvertTime(raw)

#Get metadata and display the few rows
meta = CleanMeta(raw, plate, replicate)

#Clean data
cleanraw <- CleanRaw(meta, raw, plate_time)

#Get calculations using positive controls to normalize values.
calculation = GetCalculation(raw = cleanraw, meta, sd_fold = 10)

#Formatting calculations for analysis (also compatible with graphing
#softwares used in F-SAA research)
calculation_spread = SpreadCalculation(calculation)

analysis <- GetAnalysis(calculation_spread, control = "Neg", test = "wilcox",
                        alternative = 'greater')
head(analysis)
```

GetCalculation *Perform Calculations*

#### Description

This function takes cleaned raw data and performs various analyses, including calculating the time to threshold, Rate of Amyloid Formation (RAF), Max Point Ratio (MPR), Max Slope (MS), and whether the reaction crosses the threshold (XTH).

#### Usage

```
GetCalculation(
  raw,
  meta,
  norm = FALSE,
  norm_ct,
  threshold_method = "stdv",
  time_skip = 5,
  sd_fold = 3,
  bg_fold = 3,
  rfu = 5000,
  cycle_background = 4,
  binw = 6
)
```

#### Arguments

| | |
|---|---|
| `raw` | Cleaned raw data matrix. Output from `CleanRaw()`. |
| `meta` | Cleaned meta data. Output from `CleanMeta()`. |
| `norm` | Logical. If TRUE, normalization will be performed. Default is FALSE. |
| `norm_ct` | Sample name used to normalize calculation. |
| `threshold_method` | Method for calculating threshold ('stdv', 'rfu_val', or 'bg_ratio'). |
| `time_skip` | Number of initial time points to skip when checking for threshold crossing. This helps ignore early crossings that may be due to reasons unrelated to seeding activity. |
| `sd_fold` | Fold of standard deviation to calculate the threshold for RAF (for 'stdv' method). |
| `bg_fold` | Background fold for threshold calculation (for 'bg_ratio' method). |
| `rfu` | Relative fluorescence unit values used for threshold (for 'rfu_val' method). |
| `cycle_background` | The cycle number chosen as the background for RAF and MPR calculations. |
| `binw` | Bin width for the MS calculation. |

#### Value

A data frame containing the results of the calculation.

##### Examples

```
# Define the path to the plate data file
plate_path <- system.file("extdata/20240716_p3",
                          file = '20240716_p3_plate.xlsx',
                          package = "QuICSeedR")

# Read the plate data
plate <- readxl::read_xlsx(plate_path)

# Define the path to the raw data file
raw_path <- system.file("extdata/20240716_p3",
                        file = '20240716_p3_raw.xlsx',
                        package = "QuICSeedR")
# Read the raw data
raw <- readxl::read_xlsx(raw_path)

# Get replicate data
replicate <- GetReplicate(plate)

# Ensure time displayed as decimal hours
plate_time = ConvertTime(raw)

#Get metadata and display the few rows
meta = CleanMeta(raw, plate, replicate)

#Clean data
cleanraw <- CleanRaw(meta, raw, plate_time)

#Get calculations using positive controls to normalize values.
calculation = GetCalculation(raw = cleanraw, meta, norm = TRUE, norm_ct = 'Pos')

head(calculation)
```

---

`GetReplicate` *Generate Replicate Numbers for Plate Data*

---

#### Description

This function takes a plate layout and generates a corresponding matrix of replicate numbers for each sample.

#### Usage

```
GetReplicate(plate)
```

#### Arguments

| | |
|---|---|
| `plate` | A matrix or data frame representing the plate layout, where each cell contains a sample identifier or NA for empty wells. |

#### Value

A data frame with the same dimensions as the input plate, where each cell contains the replicate number for the corresponding sample in the input plate.

#### Note

- Sample identifiers are converted to character type before processing.
- The function assumes that the input plate is organized such that replicate samples are encountered sequentially.
- The output maintains the column names from the input plate.

#### Examples

```
plate <- matrix(
  c("A", "B", "C",
    "A", "B", NA,
    "A", "C", "D"),
  nrow = 3, byrow = TRUE
)

replicates <- GetReplicate(plate)
print(replicates)
```

---

### `PlotMetric` *Create a customizable ggplot for metrics*

---

#### Description

This is wrapper function to generate a ggplot object with default options for boxplot and jittered points. Additional ggplot2 functions can be applied to the plot.

#### Usage

```
PlotMetric(
  calculation,
  x = "content",
  y = "RAF",
  fill_var = NULL,
  point = TRUE,
  box = TRUE,
  base_theme = theme_bw(),
  ...
)
```

#### Arguments

| | |
|---|---|
| `calculation` | A data frame containing the data to be plotted. Output of `GetCalculation` |
| `x` | Character string specifying the column name for the x-axis. Default is "content". |
| `y` | Character string specifying the column name for the y-axis. Default is "RAF". |
| `fill_var` | The name of the column to be used for fill color, or a vector of colors. If NULL, no fill color is applied. Default is NULL. |
| `point` | Logical. If TRUE (default), adds jittered points to the plot. |
| `box` | Logical. If TRUE (default), adds a boxplot to the plot. |
| `base_theme` | A ggplot2 theme object. Default is theme_bw(). |
| `...` | Additional ggplot2 functions to be applied to the plot. |

#### Value

A ggplot object.

#### Examples

```
# Define the path to the plate data file
plate_path <- system.file("extdata/20240716_p3",
                          file = '20240716_p3_plate.xlsx',
                          package = "QuICSeedR")

# Read the plate data
plate <- readxl::read_xlsx(plate_path)

# Define the path to the raw data file
raw_path <- system.file("extdata/20240716_p3",
                        file = '20240716_p3_raw.xlsx',
                        package = "QuICSeedR")
# Read the raw data
raw <- readxl::read_xlsx(raw_path)

# Get replicate data
replicate <- GetReplicate(plate)

# Ensure time displayed as decimal hours
plate_time = ConvertTime(raw)

#Get metadata and display the few rows
```

```
meta = CleanMeta(raw, plate, replicate)

#Clean data
cleanraw <- CleanRaw(meta, raw, plate_time)

#Get calculations using positive controls to normalize values.
calculation = GetCalculation(raw = cleanraw, meta, norm = TRUE, norm_ct = 'Pos')

#Default plot
PlotMetric(calculation)
```

---

PlotPlate *Plot Time Series Data*

---

#### Description

This function creates a faceted plot of time series data for each well in a plate layout.

#### Usage

```
PlotPlate(raw, plate_time, format = 96, f_size = 5, fill = FALSE)
```

#### Arguments

| | |
|---|---|
| raw | A data frame containing the raw plate data. The first row and first two columns are assumed to be metadata and are removed. |
| plate_time | Output from ConvertTime(). |
| format | Format of plates used in the experiment. 96 or 384. |
| f_size | font size for subtitles. |
| fill | Logical, whether to fill in missing wells with 0. Default is FALSE. |

#### Value

A ggplot object representing the plate data.

#### Examples

```
# Define the path to the plate data file
plate_path <- system.file("extdata/20240716_p3",
                          file = '20240716_p3_plate.xlsx',
                          package = "QuICSeedR")

# Read the plate data
plate <- readxl::read_xlsx(plate_path)

# Define the path to the raw data file
raw_path <- system.file("extdata/20240716_p3",
                        file = '20240716_p3_raw.xlsx',
                        package = "QuICSeedR")
# Read the raw data
raw <- readxl::read_xlsx(raw_path)
```

```
# Ensure time displayed as decimal hours
plate_time = ConvertTime(raw)

#Plot time series data for each well in a plate layout
PlotPlate(raw = raw, plate_time = plate_time, fill = TRUE)
```

---

PlotRawMulti *Plot Multiple Samples from the Cleaned Raw Data*

---

#### Description

The function visualizes the fluorescence over time for selected samples.

#### Usage

```
PlotRawMulti(
  raw,
  samples,
  legend_position = "topleft",
  xlim = NULL,
  ylim = NULL,
  custom_colors = NULL,
  xlab = "Time (h)",
  ylab = "Fluorescence",
  linetypes = NULL
)
```

#### Arguments

| | |
|---|---|
| `raw` | Cleaned raw data. Output from `GetCleanRaw`. |
| `samples` | The names of the samples to plot. |
| `legend_position` | Position of legend. Default is "topleft". Choose from "bottomright", "bottom", "bottomleft", "left", "topleft", "top", "topright", "right" and "center". |
| `xlim` | Numeric vector giving the x coordinate range (hours). If NULL (default), limits are computed from the data. |
| `ylim` | Numeric vector giving the y coordinate range (relative fluorescence units). If NULL (default), limits are computed from the data. |
| `custom_colors` | An optional vector of colors to be used for plotting. If NULL (default), the function will use the original color scheme. |
| `xlab` | Label for x-axis. |
| `ylab` | Label for y-axis. |
| `linetypes` | Vector of line types to use for each line. Can be integer codes (1:6) or character codes ("solid", "dashed", "dotted", "dotdash", "longdash", "twodash"). All lines will be solid by default. |

#### Value

A plot displaying the fluorescence of the selected samples over time.

#### Examples

```
# Define the path to the plate data file
plate_path <- system.file("extdata/20240716_p3",
                          file = '20240716_p3_plate.xlsx',
                          package = "QuICSeedR")

# Read the plate data
plate <- readxl::read_xlsx(plate_path)

# Define the path to the raw data file
raw_path <- system.file("extdata/20240716_p3",
                        file = '20240716_p3_raw.xlsx',
                        package = "QuICSeedR")
# Read the raw data
raw <- readxl::read_xlsx(raw_path)

# Get replicate data
replicate <- GetReplicate(plate)

# Ensure time displayed as decimal hours
plate_time = ConvertTime(raw)

#Get metadata and display the few rows
meta = CleanMeta(raw, plate, replicate)

#Clean data
cleanraw <- CleanRaw(meta, raw, plate_time)

#Plot fluorescence curves from negative and positive controls
PlotRawMulti(cleanraw, c("Neg", "Pos"))
```

---

### `PlotRawSingle` *Plot a Single Sample from the Cleaned Raw Data*

---

#### Description

The function plots fluorescence for selected sample over time.

#### Usage

```
PlotRawSingle(
  raw,
  sample,
  legend_position = "topleft",
  xlim = NULL,
  ylim = NULL,
  custom_colors = NULL,
  xlab = "Time (h)",
  ylab = "Fluorescence",
  linetypes = NULL
)
```

##### Arguments

- `raw` Cleaned raw data. Output from `GetCleanRaw`
- `sample` The name of the sample to plot.
- `legend_position` Position of legend. Default is "topleft".
- `xlim` Numeric vector giving the x coordinate range (hours). If NULL (default), limits are computed from the data. Choose from "bottomright", "bottom", "bottomleft", "left", "topleft", "top", "topright", "right" and "center".
- `ylim` Numeric vector giving the y coordinate range (relative fluorescence units). If NULL (default), limits are computed from the data.
- `custom_colors` An optional vector of colors to be used for plotting. If NULL (default), the function will use the original color scheme.
- `xlab` Label for x-axis.
- `ylab` Label for y-axis.
- `linetypes` Vector of line types to use for each line. Can be integer codes (1:6) or character codes ("solid", "dashed", "dotted", "dotdash", "longdash", "twodash"). All lines will be solid by default.

##### Value

A plot displaying the fluorescence of the selected sample over time.

##### Examples

```
# Define the path to the plate data file
plate_path <- system.file("extdata/20240716_p3",
                          file = '20240716_p3_plate.xlsx',
                          package = "QuICSeedR")

# Read the plate data
plate <- readxl::read_xlsx(plate_path)

# Define the path to the raw data file
raw_path <- system.file("extdata/20240716_p3",
                        file = '20240716_p3_raw.xlsx',
                        package = "QuICSeedR")
# Read the raw data
raw <- readxl::read_xlsx(raw_path)

# Get replicate data
replicate <- GetReplicate(plate)

# Ensure time displayed as decimal hours
plate_time = ConvertTime(raw)

#Get metadata and display the few rows
meta = CleanMeta(raw, plate, replicate)

#Clean data
cleanraw <- CleanRaw(meta, raw, plate_time)

#Plot fluorescence curves from positive controls
```

```
PlotRawSingle(cleanraw, "Pos")
```

---

#### SpreadCalculation *Spread Calculation Data*

---

##### Description

This function takes a data frame containing metadata and calculation results, and spreads the results into a list of data frames for selected calculation term.

##### Usage

```
SpreadCalculation(
  calculation,
  id_col = "content",
  rep_col = "replicate",
  terms = c("RAF", "MPR", "MS")
)
```

##### Arguments

| | |
|---|---|
| calculation | A data frame containing the metadata and results of the calculation. Output from GetCalculation(). |
| id_col | The name of the column in calculation that identifies the content. Default is 'content'. |
| rep_col | The name of the column in calculation that identifies the replicate. Default is 'replicate'. |
| terms | A vector of column names to spread. Defaults to 'RAF', 'MPR', and 'MS'. |

##### Value

A list of data frames containing the spread results of the calculation. Each data frame is compatible with various graphing software, particularly GraphPad Prism, which is the most commonly used graphing tool in F-SAA research.

##### Examples

```
calculation <- data.frame(
  content = rep(c("A", "B", "C"), each = 2),
  replicate = rep(1:2, 3),
  time_to_threshold = rnorm(6),
  RAF = rnorm(6),
  MPR = rnorm(6),
  MS = rnorm(6)
  )

calculation_spread = SpreadCalculation(calculation)
print(calculation_spread)
```

`SummarizeResult` *Summarize Analysis Results*

#### Description

This function combines analysis results from multiple tests with metadata, and determines overall significance based on a specified method. By default, it evaluates sample-level result by calculating the percentage of technical replicates that exceed the pre-defined threshold.

#### Usage

```
SummarizeResult(
  analysis = NULL,
  calculation,
  sig_method = "xth_percent",
  method_threshold = 50
)
```

#### Arguments

| | |
|---|---|
| `analysis` | Output of `GetAnalysis()`. |
| `calculation` | Output of `GetCalculation()`. |
| `sig_method` | Specifies the approach for determining sample-level result. Available options include "xth_percent", "metric_count", "xth_count", or any metric name present in the analysis list. The default is "xth_percent". |
| `method_threshold` | Defines the threshold value for the "metric_count", "xth_count", and "xth_percent" methods. This parameter defaults to 50. |

#### Value

A data frame summarizing results of the analysis and calculation, with columns:

- result: Overall result based on the chosen method
- position: Location of the sample replications based on plate layout
- method: Method chosen for determination of the result
- *_sig: Significance indicator for each metric
- *_p: P-value or adjusted p-value for each metric
- metric_count: Count of significant results across all analyses
- xth_count: The number of reactions crossing the fluorescent threshold
- total_rep: Total number of replicates
- xth_percent: Percentage of replicates crossing the fluorescent threshold

#### Examples

```
# Define the path to the plate data file
plate_path <- system.file("extdata/20240716_p3",
                          file = '20240716_p3_plate.xlsx',
                          package = "QuICSeedR")

# Read the plate data
plate <- readxl::read_xlsx(plate_path)

# Define the path to the raw data file
raw_path <- system.file("extdata/20240716_p3",
                        file = '20240716_p3_raw.xlsx',
                        package = "QuICSeedR")
# Read the raw data
raw <- readxl::read_xlsx(raw_path)

# Get replicate data
replicate <- GetReplicate(plate)

# Ensure time displayed as decimal hours
plate_time = ConvertTime(raw)

#Get metadata and display the few rows
meta = CleanMeta(raw, plate, replicate)

#Clean data
cleanraw <- CleanRaw(meta, raw, plate_time)

#Get calculations using positive controls to normalize values.
calculation = GetCalculation(raw = cleanraw, meta, sd_fold = 10)

#Formatting calculations for analysis (also compatible with graphing softwares used in F-SAA
#research)
calculation_spread = SpreadCalculation(calculation)

#Get analysis comparing samples to negative control using the one-tailed Wilcox Rank-Sum test.
analysis <- GetAnalysis(calculation_spread, control = "Neg", test = "wilcox",
                        alternative = 'greater')

#Summarization of results. Default method is rate of amyloid formation.
result <- SummarizeResult(analysis, calculation)

head(result)
```

### Index

BulkProcessing, 2
BulkReadMARS, 3

CleanMeta, 4
CleanRaw, 5
ConvertTime, 6

GetAnalysis, 7
GetCalculation, 9
GetReplicate, 10

PlotMetric, 11
PlotPlate, 13
PlotRawMulti, 14
PlotRawSingle, 15

SpreadCalculation, 17
SummarizeResult, 18
