## Supplementary material for "QuICSeedR: An R package for analyzing fluorophore-assisted seed amplification assay data": Basic-Tutorial.pdf

### QuICSeedR Basic Tutorial

Manci Li, PhD

2024-08-27

#### Introduction

This tutorial provides a basic guide to using the QuICSeedR package for analyzing data from fluorophore-assisted seed amplification assays (F-SAAs). We'll focus on the basic workflow and flexible plate format feature by demonstrating its application with environmental soil samples from Chronic Wasting Disease (CWD) positive-sites. For details of experiments, contact Dr. Stuart Lichtenberg at.

#### 1. Installation and Setup

You can install the development version of QuICSeedR from GitHub with:

```r
# install.packages("devtools")
# devtools::install_github("mancili/QuICSeedR")
```

Load libraries before analysis

```r
library(tidyverse)
library(QuICSeedR)
library(knitr)
library(readxl)
```

In this section, I demonstrate the workflow's compatibility with both 96- and 384-well plate formats, providing a detailed explanation of the functions involved. The package predominantly features automatic recognition of plate format, with the exception of `PlotPlate()` function. The design ensures computability of experimental data while maintaining flexibility across different experimental setups.

**2. Workflow**

```
#Load data from sample 96-well plate
plate_96 = read_xlsx('data/20240716_s1_plate.xlsx')
raw_96 = read_xlsx('data/20240716_s1_raw.xlsx')

#Load data from sample 384-well plate
plate_384 = read_xlsx('data/20230808_M12_plate.xlsx')
raw_384 = read_xlsx('data/20230808_M12_raw.xlsx')
```

**1) Load sample data and plot plate overview** BMG readers have a different time format. To consistently get decimal hours, you can run:

```
#96-well plate
plate_time_96 = ConvertTime(raw_96)

#384-well plate
plate_time_384 = ConvertTime(raw_384)
```

The fluorescence can be viewed with `PlotPlate()`, default plate format is 96-well. Additionally, `PlotPlate()` supports partial plate visualization when the `fill` parameter is set to `TRUE` (default is `FALSE`).

```
#96-well plate
PlotPlate(raw_96, plate_time_96)
```

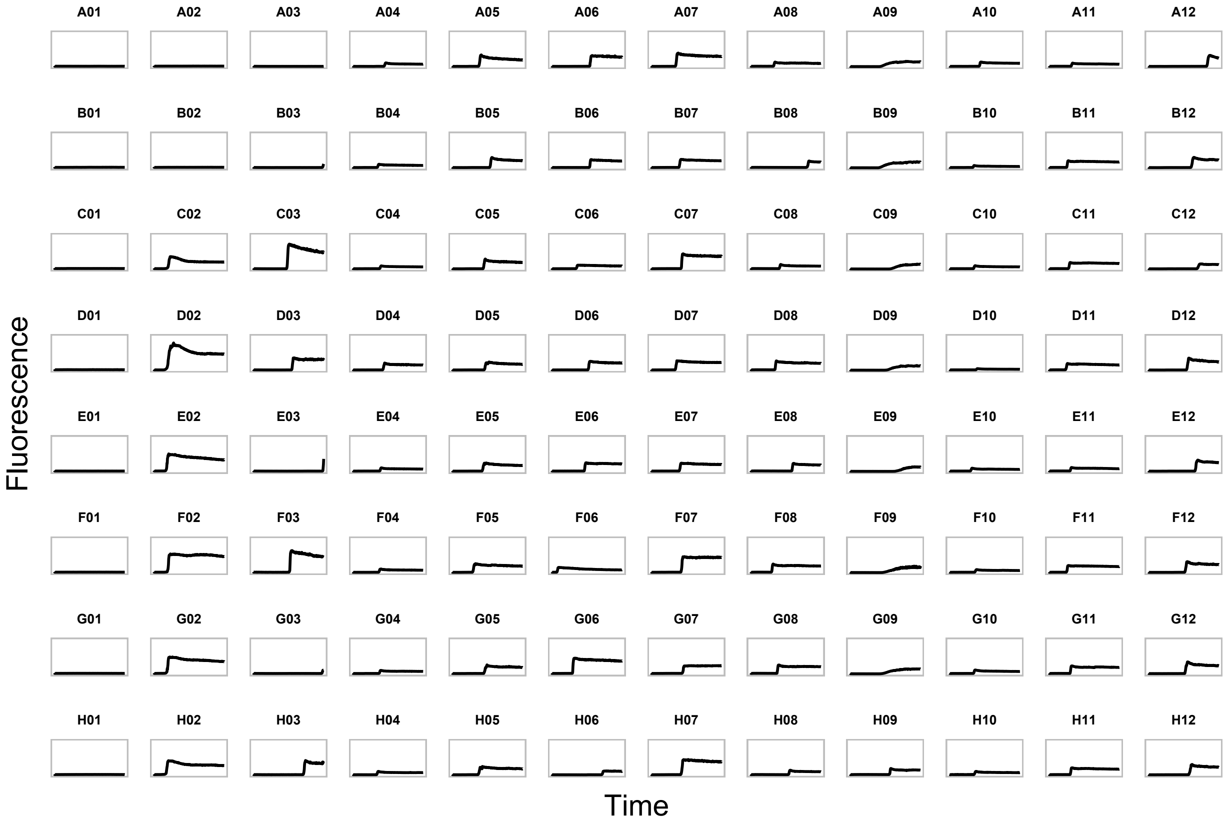

Although the code is provided here, 384-well plots are easier to view interactively using the `plotly` package.

```
#install.packages('plotly')
library(plotly)

p = PlotPlate(raw_384, plate_time_384, format = 384)
interactive_plot <- ggplotly(p)
interactive_plot
```

**2) Data cleaning** In addition to `ConvertTime()`, the data cleaning process is primarily executed through three key functions—`GetReplicate()`, `CleanMeta()`, and `CleanRaw()`, each addressing specific aspects of data preparation and organization.

To accommodate different plate layout without manual entry of technical replicate, `GetReplicate()` function is used to assign the replicate information.

```
#96-well plate
replicate_96 = GetReplicate(plate_96)

#384-well plate
replicate_384 = GetReplicate(plate_384)
```

The `CleanMeta()` function generates experimental metadata, offering flexible options for content manipulation and data cleaning. It allows users to split sample names into multiple columns using the `split_content` parameter (default FALSE), with the splitting delimiter specified by `split_by` (default "_"). When splitting is enabled, users must provide column names for the resulting split data via the split_into parameter. The function also includes an option to remove rows containing NA values through the `del_na` parameter (default TRUE), facilitating dataset refinement. Additional metadata can be added onto the output of `CleanMeta()` without impacting downstream analysis. These features enable efficient preprocessing of metadata, enhancing its structure for subsequent analysis and integration with experimental results.

```
#96-well plate
meta_96 = CleanMeta(raw_96, plate_96, replicate_96)

#384-well plate
meta_384 = CleanMeta(raw_384, plate_384, replicate_384)
```

The `CleanRaw()` function is primarily designed to process MARS output, reflecting its current status as the standard instrument for measuring thioflavin T (ThT) fluorescence in F-SAAs. The function generates a cleaned matrix where column names represent sample names and replicate numbers, while row names indicate time in decimal hours. Although optimized for MARS output, `CleanRaw()` is adaptable to potential future instrumentation or data formats. Users can utilize the function's downstream analysis capabilities by converting their data to the required format.

```
#96-well plate
cleanraw_96 = CleanRaw(meta_96, raw_96, plate_time_96)

#384-well plate
cleanraw_384 = CleanRaw(meta_384, raw_384, plate_time_384)
```

Output of `CleanRaw()` can be used to visualize sample of interest using `PlotRawMulti()` or `PlotRawSingle()` functions. For example, to check positive ("Pos" on 96-well plate; "pos" on 384-well plate) and negative ("Neg") controls of the plate:

```r
par(mfrow = c(1,2))
#96-well plate
PlotRawMulti(cleanraw_96, c("Neg", "Pos"))

#384-well plate
PlotRawSingle(cleanraw_384, "pos")
```

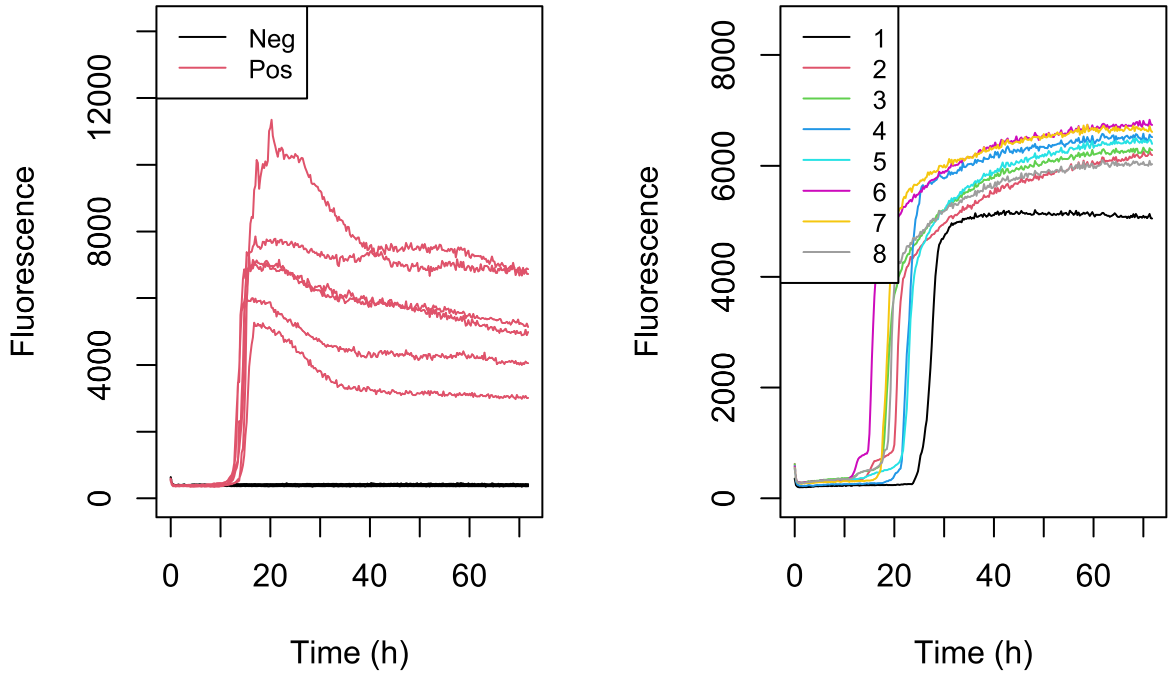


```r
par(mfrow = c(1,1))
```

**3) Calculation** The `GetCalculation()` function is a core component of the QuICSeedR package, designed to perform various analyses on cleaned raw data from seed amplification assays. It calculates several key metrics that are crucial for interpreting the results of these experiments:

- Time to Threshold: The time point at which the fluorescence signal crosses a defined threshold. `cycle_background` specifies which cycle to use as the reading for the background reading. You can choose between standard deviation ("stdv") or background ratio ("bg_ratio") methods for threshold calculation. `sd_fold` and `bg_fold` are parameters for fine-tuning threshold calculations (how many times of the chosen background cycle or its mean) based on the chosen method. If a reaction never crosses threshold, NA will be designated.

- Threshold Crossing (XTH): A binary indicator of whether the reaction crosses the threshold. Reactions crossing the threshold receive 1 while those that don't receive 0.

- Rate of Amyloid Formation (RAF): The rate at which amyloid aggregates form during the assay, calculate as the inverse of time to threshold. `time_skip` helps ignore early threshold crossings that might be due to experimental artifacts. If a reaction never crosses threshold, 0 will be designated.

- Max-point Ratio (MPR): The ratio of the maximum fluorescence to the background fluorescence.
- Max Slope (MS): The maximum rate of change in fluorescence over time. `binw` can be used to define the bin width (time window) for smoothing the slope.

Furthermore, the function offers a data normalization option through the `norm` parameter, with `norm_ct` specifying the reference sample for normalization. This normalization feature is particularly crucial when utilizing the RAF as a semi-quantitative measure of misfolded protein concentration, as it enhances the comparability and reliability of results across different experimental instruments, runs, and conditions.

```
#96-well plate
calculation_96 = GetCalculation(cleanraw_96, meta_96, norm = TRUE, norm_ct = 'Pos')

#384-well plate
calculation_384 = GetCalculation(cleanraw_384, meta_384, norm = TRUE, norm_ct = 'pos')
```

Calculated metrics of interest can be visualized through `PlotMetric()`.

```
#384-well plate
PlotMetric(calculation_384, y = "MS")
```

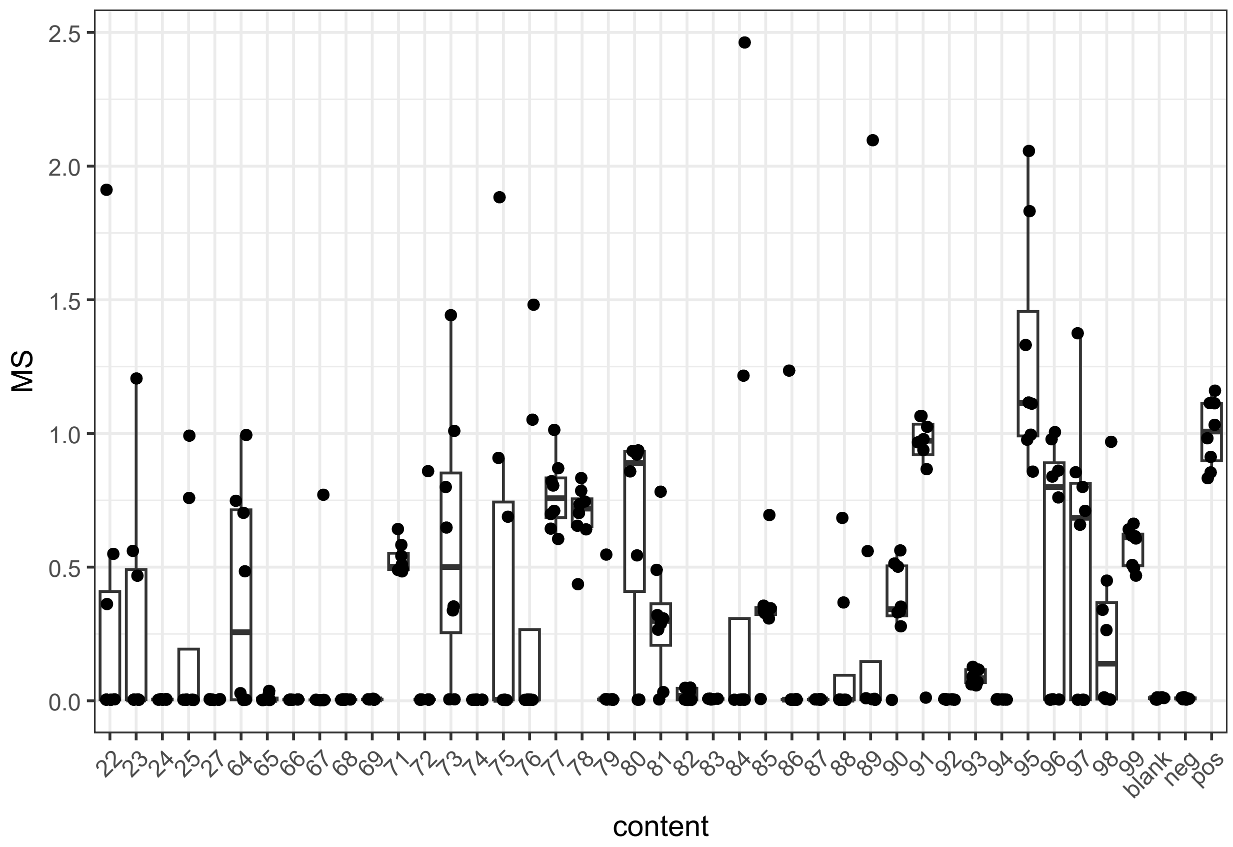


This is also compatible with parameters in the ggplot package.

```
PlotMetric(calculation_96, y = "MS", point = FALSE, box = FALSE,
  boxplot= geom_boxplot(color = 'gray'),
  scatter = geom_point(color = "blue") ,
```

```
  xlab = xlab("Sample ID"),
  ylab =ylab("Normalized Max Slope"))
```

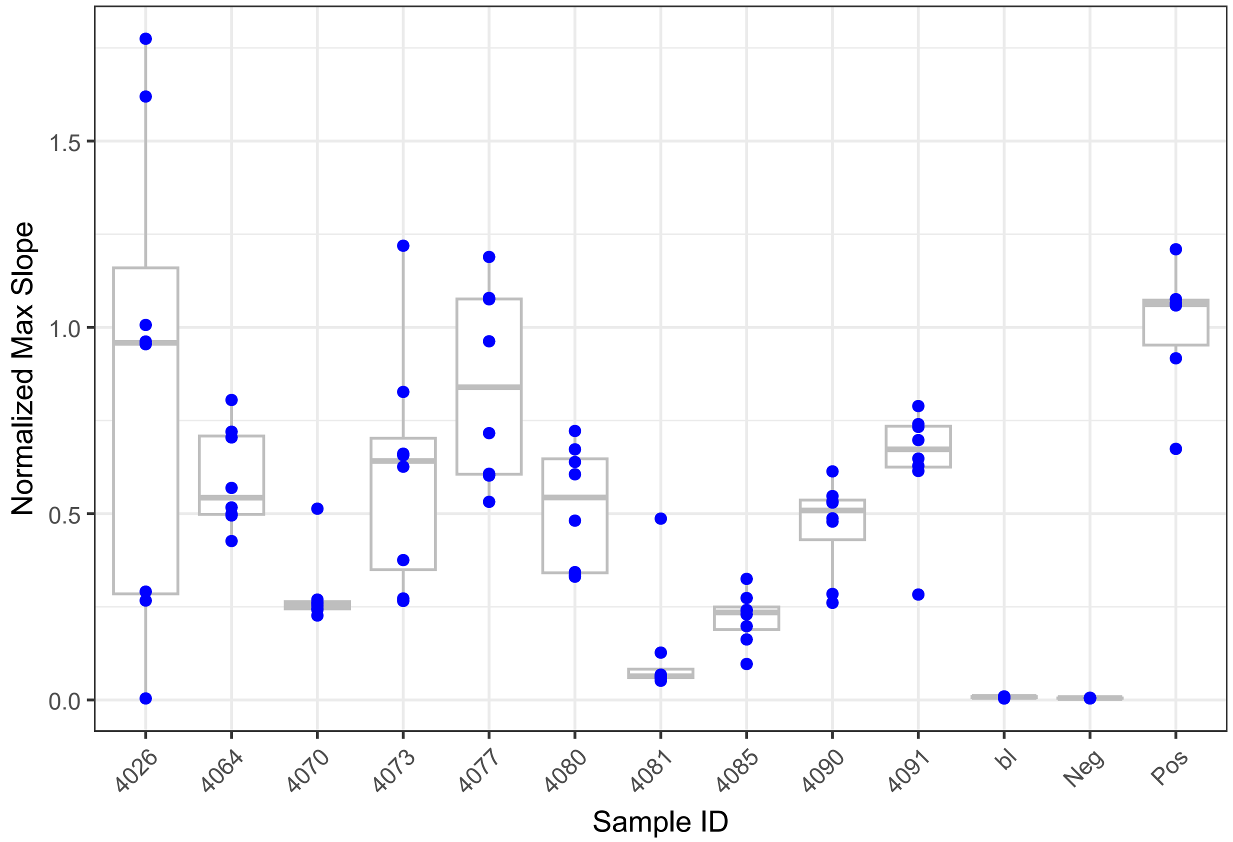


To obtain files suitable for plotting in the GraphPad Prism software, use `SpreadCalculation()`. The function by default performs analysis on all three metrics: RAF, MS, and MPR. However, there is flexibility to analyze any combination of these metrics (one, any two, or all three metrics). The selection determines which metrics undergo statistical analysis.

Here, I am interested in RAF. See references article for details.

```
calculation_spread_96 = SpreadCalculation(calculation_96, terms = 'RAF')

#view
kable(round(calculation_spread_96[['RAF']], 2))
```

| bl | Neg | 4026 | 4070 | 4064 | 4073 | 4077 | 4080 | 4081 | 4085 | 4090 | 4091 | Pos |
|---|---|---|---|---|---|---|---|---|---|---|---|---|
| 0.00 | 0.00 | 0.00 | 0.36 | 0.43 | 0.30 | 0.47 | 0.49 | 0.36 | 0.38 | 0.51 | 0.20 | 0.91 |
| NA | 0.29 | 0.16 | 0.42 | 0.38 | 0.35 | 0.40 | 0.28 | 0.24 | 0.53 | 0.53 | 0.24 | 0.99 |
| 0.24 | 0.00 | 0.16 | 0.46 | 0.30 | 0.30 | 0.41 | 0.20 | 0.37 | 0.48 | 0.64 | 0.27 | 1.01 |
| NA | 0.20 | 0.32 | 0.43 | 0.55 | 2.11 | 0.38 | 0.54 | 0.33 | 0.45 | 0.65 | 0.31 | 1.22 |
| 0.00 | 0.00 | 0.34 | 0.42 | 0.37 | 0.46 | 0.38 | 0.39 | 0.27 | 0.47 | 0.58 | 0.23 | 0.97 |
| 0.86 | 0.00 | 0.30 | 0.38 | 0.35 | 0.31 | 0.47 | 0.47 | 0.30 | 0.43 | 0.68 | 0.30 | 0.89 |
| NA | NA | 0.17 | 0.43 | 0.36 | 0.55 | 0.37 | 0.43 | 0.33 | 0.47 | 0.53 | 0.31 | NA |
| NA | NA | 0.23 | 0.47 | 0.43 | 0.23 | 0.38 | 0.30 | 0.29 | 0.46 | 0.53 | 0.28 | NA |

**4) Statistical Analysis** The `GetAnalysis()` function conducts statistical analysis on data processed by `SpreadCalculation()`. It compares samples against a control (typically a negative control) using one of three statistical tests: Wilcoxon rank-sum test ("wilcox", default), Yuen's test for trimmed means ("yuen"), Student's t-test ("t-test"). The Wilcoxon test is non-parametric and suitable for non-normally distributed data and the t-test assumes normality and is best for continuous data. While Yuen's test is robust against outliers and non-normality, minimum of 8 replicates is recommended as the trim level is set at 10%. Consult the documentation for each statistical test to ensure appropriate application based on your data's characteristics and distribution.

The function allows specification of the alternative hypothesis (`alternative`) as two-sided ("two.sided", default), less than ("less"), or greater than ("greater"). While p-value adjustment (`adjust_p`) for multiple comparisons using Benjamini-Hochberg is available, it's disabled by default to maintain sensitivity, as controlling the false discovery rate might reduce the ability to detect true positives. The significance level can be set using the `alpha` parameter, with a default of 0.05. The processes for 96-well and 384-well plates are identical for the following steps, so we'll demonstrate using the 96-well plate format as an example.

Here, I am using the one-tailed Wilcoxon Rank-Sum test for RAF.

```r
#96-well plate
analysis_96 = GetAnalysis(calculation_spread = calculation_spread_96,
                          control = 'Neg',
                          test = 'wilcox',
                          alternative = 'greater',
                          alpha = 0.05)
kable(analysis_96$RAF)
```

| | statistic | p_value | significant |
|---|---|---|---|
| bl | 15.0 | 0.27409 | |
| Neg | 18.0 | 0.53814 | |
| 4026 | 37.0 | 0.04943 | * |
| 4070 | 48.0 | 0.00108 | ** |
| 4064 | 48.0 | 0.00108 | ** |
| 4073 | 47.0 | 0.00166 | ** |
| 4077 | 48.0 | 0.00107 | ** |
| 4080 | 46.0 | 0.00250 | ** |
| 4081 | 45.5 | 0.00303 | ** |
| 4085 | 48.0 | 0.00107 | ** |
| 4090 | 48.0 | 0.00107 | ** |
| 4091 | 42.0 | 0.01110 | * |
| Pos | 36.0 | 0.00217 | ** |

**5) Summary of analysis** `SummarizeResult()` can be used to see the summary of the analysis. I am using the results of RAF analysis to determine whether sample had seeding activity.

```
#96-well plate
kable(SummarizeResult(analysis = analysis_96,
                      calculation = calculation_96,
                      sig_method = 'RAF'))
```

| content | result | method | position | RAF_sig | RAF_p | metric_count | xth_count | total_rep | xth_percent |
|---|---|---|---|---|---|---|---|---|---|
| bl | | RAF | A01-B01-C01-D01 | | 0.27409 | 0 | 2 | 4 | 50.00 |
| Neg | | RAF | A02-B02-E01-F01-G01-H01 | | 0.53814 | 0 | 2 | 6 | 33.33 |
| 4026 | * | RAF | A03-B03-C03-D03-E03-F03-G03-H03 | * | 0.04943 | 1 | 7 | 8 | 87.50 |
| 4070 | * | RAF | A04-B04-C04-D04-E04-F04-G04-H04 | ** | 0.00108 | 1 | 8 | 8 | 100.00 |
| 4064 | * | RAF | A05-B05-C05-D05-E05-F05-G05-H05 | ** | 0.00108 | 1 | 8 | 8 | 100.00 |
| 4073 | * | RAF | A06-B06-C06-D06-E06-F06-G06-H06 | ** | 0.00166 | 1 | 8 | 8 | 100.00 |
| 4077 | * | RAF | A07-B07-C07-D07-E07-F07-G07-H07 | ** | 0.00107 | 1 | 8 | 8 | 100.00 |
| 4080 | * | RAF | A08-B08-C08-D08-E08-F08-G08-H08 | ** | 0.00250 | 1 | 8 | 8 | 100.00 |
| 4081 | * | RAF | A09-B09-C09-D09-E09-F09-G09-H09 | ** | 0.00303 | 1 | 8 | 8 | 100.00 |
| 4085 | * | RAF | A10-B10-C10-D10-E10-F10-G10-H10 | ** | 0.00107 | 1 | 8 | 8 | 100.00 |
| 4090 | * | RAF | A11-B11-C11-D11-E11-F11-G11-H11 | ** | 0.00107 | 1 | 8 | 8 | 100.00 |
| 4091 | * | RAF | A12-B12-C12-D12-E12-F12-G12-H12 | * | 0.01110 | 1 | 8 | 8 | 100.00 |
| Pos | * | RAF | C02-D02-E02-F02-G02-H02 | ** | 0.00217 | 1 | 6 | 6 | 100.00 |
