## Supplementary material for "QuICSeedR: An R package for analyzing fluorophore-assisted seed amplification assay data": ParallelProcessing.R

```
#' Parallel Processing of Multiple Experiments
#'
#' This function processes multiple experiments in parallel, performing a series of operations
#' including time conversion, metadata cleaning, raw data cleaning, calculations, analysis,
#' and result summarization.
#'
#' @param data Compiled data of experiments.  
#'   
#' @param params A list of parameters for various processing steps. If a parameter is not
#'   provided, default values will be used. `control` used in `GetAnalysis()` is required.
#'   parameters can be provided for the following functions:
#'   \itemize{
#'     \item `ConvertTime`
#'     \item `CleanMeta`
#'     \item `CleanRaw`
#'     \item `GetCalculation`
#'     \item `SpreadCalculation`
#'     \item `GetAnalysis`
#'     \item `SummarizeResult`
#'   }
#'
#' @return A list containing three elements:
#'   \itemize{
#'     \item combined_calculation: A data frame of combined calculations from all experiments
#'     \item combined_cleanraw: Cleaned raw data for each experiment
#'     \item combined_result: A data frame of combined results from all experiments
#'   }
#'
#' @examples
#' dontrun{
#' 
#' #Set parameters that will be used in the process
#' 
#' myparams = list(
#' CleanMeta = list(split_content = TRUE, split_into = c("dilution", "sampleID")),
#' GetAnalysis = list(control = 'neg', alternative = "greater"),
#' SummarizeResult = list(sig_method = 'metric_count', method_threshold = 3)
#' )
#' 
#' results = ParallelProcessing(data = my_data, params = my_params)
#' 
#' # Access combined results
#' combined_results <- results$combined_result
#'
#' # Access cleaned raw data for a specific plate
#' cleaned_raw_data_plate1 <- results$combined_cleanraw[[1]]
#'
#' @export
ParallelProcessing = function(data, params = list()) { 
  
  subcalculation <- list()
  subcleanraw <- list()
  subresult <- list()
  
  for (j in 1:length(data)) {
    plate <- data[[j]]$plate
    raw <- data[[j]]$raw
    replicate <- data[[j]]$replicate
    
    plate_time <- do.call(ConvertTime, c(list(raw), params$ConvertTime %||% list()))
    
    meta <- do.call(CleanMeta, c(list(raw = raw, plate = plate, replicate = replicate), 
                                 params$CleanMeta %||% list()))
    
    raw <- do.call(CleanRaw, c(list(meta = meta, raw = raw, plate_time = plate_time), 
                               params$CleanRaw %||% list()))
    
    calculation <- do.call(GetCalculation, c(list(raw = raw, meta = meta), 
                                             params$GetCalculation %||% list()))
    
    calculation_spread <- do.call(SpreadCalculation, c(list(calculation), 
                                                       params$SpreadCalculation %||% list()))
    
    analysis <- do.call(GetAnalysis, c(list(calculation_spread), 
                                       params$GetAnalysis %||% list()))
    
    result <- do.call(SummarizeResult, c(list(analysis = analysis, calculation = calculation), 
                                         params$SummarizeResult %||% list()))
    
    subcalculation[[j]] <- calculation
    subcleanraw[[j]] <- raw
    subresult[[j]] <- result
  }
  
  names(subcalculation) = names(data)
  names(subcleanraw) = names(data)
  names(subresult) = names(data)
  
  subresult <- lapply(names(subresult), function(name) {
    data <- subresult[[name]]
    data$plate_name <- name
    data
  })
  fullresult = do.call(rbind, subresult)
  
  subcalculation <- lapply(names(subcalculation), function(name) {
    data <- subcalculation[[name]]
    data$plate_name <- name
    data
  })
  fullcalc = do.call(rbind, subcalculation)
  
  return(list(combined_calculation = fullcalc,
              combined_cleanraw = subcleanraw,
              combined_result = fullresult))
}
```
