## Supplementary material for "QuICSeedR: An R package for analyzing fluorophore-assisted seed amplification assay data": saveRDS.R

```
flip_and_replace <- function(x) {
  process_single <- function(match) {
    parts <- strsplit(match, "\\^")[[1]]
    base <- parts[1]
    exponent <- parts[2]
    
    if (exponent == "-1") {
      return(paste0("10e1_", base))
    } else if (exponent == "-2") {
      return(paste0("10e2_", base))
    } else {
      return(paste0("10e3_", base))
    } 
  }
  
  pattern <- "\\w+\\^-[123]\\b"
  matches <- gregexpr(pattern, x)
  regmatches(x, matches) <- lapply(regmatches(x, matches), function(match) sapply(match, process_single))
  
  return(x)
}


library(stringr)
library(readxl)
library(tidyverse)
library(QuICAnalysis)

folders <- list.dirs(path = ".", recursive = FALSE)

mylist = vector(mode = 'list', length = length(folders))

listnames = sub("\\./", "", folders)
names(mylist) = c(listnames)


for (i in 1:length(folders)) {
  folder <- folders[i]
  
  files <- list.files(path = folder, pattern = "*.xlsx", full.names = TRUE)
  
  plate_path <- files[grepl("[Pp]late", files)]
  raw_path <- files[grepl("[Rr]aw", files)]
  
  plate_data <- read_xlsx(plate_path)
  raw_data <- read_xlsx(raw_path)
  replicate_data <- GetReplicate(plate_data)
 
  mylist[[i]][['plate']] <- plate_data
  mylist[[i]][["plate"]] <- data.frame(lapply(mylist[[i]][["plate"]], flip_and_replace))
  
  mylist[[i]][['raw']] <- raw_data
  mylist[[i]][['replicate']] <- replicate_data
}


#saveRDS(mylist, file = 'multiplate_grinderpilot.rds')
```
