## Supplementary material for "QuICSeedR: An R package for analyzing fluorophore-assisted seed amplification assay data": script.R

```
flip_and_replace <- function(x) {
  process_single <- function(match) {
    parts <- strsplit(match, "\\^")[[1]]
    base <- parts[1]
    exponent <- parts[2]
    
    if (exponent == "-1") {
      return(paste0("10e1_", base))
    } else if (exponent == "-2") {
      return(paste0("10e2_", base))
    } else {
      return(paste0("10e3_", base))
    } 
  }
  
  pattern <- "\\w+\\^-[123]\\b"
  matches <- gregexpr(pattern, x)
  regmatches(x, matches) <- lapply(regmatches(x, matches), function(match) sapply(match, process_single))
  
  return(x)
}

library(stringr)
library(tidyverse)
library(QuICSeedR)
library(gplots)
library(viridis)
library(readxl)

setwd("~/R/QuICAnalysis_Manuscript/pdf_vignettes")

grinder_list = BulkReadMARS('./data/grinder', 
                           plate_subfix = 'plate',
                           raw_subfix = 'raw',
                           helper_func = flip_and_replace)
params = list(
  CleanMeta = list(split_content = TRUE, 
                   split_into = c("dilution", "sampleID")),
  GetCalculation = list(norm = TRUE, norm_ct = 'pos', 
                        sd_fold = 10,
                        cycle_background = 6), #fix cycle background in 
  GetAnalysis = list(control = 'neg', alternative = "greater"),
  SummarizeResult = list(sig_method = 'metric_count', method_threshold = 3)
)

results = BulkProcessing(data = grinder_list, params = params)


#'10e1_21P'
PlotRawMulti(results$combined_cleanraw$`20221114_M2`, c('10e1_21P', 'neg'))


#Extract calculation of metrics and prepare the data for adding study metadata
result = results$combined_calculation
sel = c('blank', 'pos', 'neg')
sel = which(result$content %in% sel)
result = result[-sel, ]
colnames(result)[7] = 'SampleID'

#Add study metadata
metadata = read_xlsx('./data/grinder_summary.xlsx')
full_result = right_join(metadata, result)

#prepare for visualization
sel0 = which(full_result$Treatment == 'none')
sel1 = which(full_result$CWD == 'negative')
data = full_result[c(sel0,sel1),]

sel2 = which(results$combined_result$result == '*') 
result_sel = results$combined_result[sel2,]

data = inner_join(data, result_sel)
data = rbind( full_result[1:36,colnames(data)[1:19]], data[,1:19])
data$content = factor(data$content, levels = rev(unique(data$content)))


#show 
col_palette <- viridis(50, option = 'turbo') 

ggplot(data, aes(x = replicate, y = content, fill = RAF)) + 
  geom_tile(color = "white",
            lwd = 1,
            linetype = 1) +
  scale_y_discrete(limits = rev(levels(factor(data$content))))+
  scale_fill_gradientn(colors = col_palette) +
  theme_bw() +
  coord_flip() +
  theme(axis.text.x = element_text(angle = 45, hjust = 1),
        aspect.ratio = 1/5)


png(filename = 'grinder_RAF.png', width = 3, height = 5, units = 'in', res = 300)
ggplot(data, aes(x = replicate, y = content, fill = RAF)) + #applies to MPR and MS as well
  geom_tile(color = "white",
            lwd = 1,
            linetype = 1) +
  coord_fixed() +
  scale_fill_gradientn(colors = col_palette) +
  theme_bw() 
dev.off()
  
#not show 
png(filename = 'grinder_MPR.png', width = 3, height = 5, units = 'in', res = 300)
ggplot(data, aes(x = replicate, y = content, fill = MPR)) +
  geom_tile(color = "white",
            lwd = 1,
            linetype = 1) +
  coord_fixed() +
  scale_fill_gradientn(colors = col_palette) +
  theme_bw()
dev.off()

png(filename = 'grinder_MS.png', width = 3, height = 5, units = 'in', res = 300)
ggplot(data, aes(x = replicate, y = content, fill = MS)) +
  geom_tile(color = "white",
            lwd = 1,
            linetype = 1) +
  coord_fixed() +
  scale_fill_gradientn(colors = col_palette) +
  theme_bw()
dev.off()
```
