## Supplementary material for "QuICSeedR: An R package for analyzing fluorophore-assisted seed amplification assay data": Versatile-Application.pdf

### Versatile Usage of QuICSeedR

Manci Li, PhD

2024-08-27

#### 1. Introduction

This document demonstrates how QuICSeedR can achieve enhanced analytical capabilities, scalable high-throughput processing with `BulkReadMARS` and `BulkProcessing`, and versatile applications across various sample types, neurodegenerative diseases, and fluorescence-assisted seed amplification assays (F-SAAs), including real-time quaking-induced conversion (RT-QuIC) (Wilham et al., 2010; Atarashi et al., 2011), nanoparticle-enhanced RT-QuIC (nano-QuIC) (Christenson et al., 2023), and fluorophore-assisted protein misfolding cyclic amplification (F-PMCA) (Shahnawaz et al., 2017; Singer et al., 2020, 2021). For installation, setup, and workflow instructions, refer to the basic tutorial.

The metrics used in this tutorial from `QuICSeedR` include Time to Threshold, which marks the point when the fluorescence signal crosses a defined threshold, and Threshold Crossing (XTH), a binary indicator of whether this threshold is crossed. The Rate of Amyloid Formation (RAF) measures the speed of amyloid aggregation, calculated as the inverse of the time to threshold. Max-point Ratio (MPR) is the ratio of the maximum fluorescence to the background fluorescence, while Max Slope (MS) captures the highest rate of change in fluorescence over time. These metrics help quantify the differences in amyloid formation and provide insights into the effectiveness of the assay. See the basic tutorial for details.

#### 2. Versatile Use Cases

**1) Achieving scalability by high throughput analysis** This section demonstrates the toolkit's capability to handle large-scale datasets efficiently. Its performance is showcased by using a subset of RT-QuIC data from a chronic wasting disease (CWD) environmental swab study [citation needed]. In this exemplary application, QuICSeedR successfully:

- Read in, processed, and analyzed 242 samples with automation
- Managed 1,152 individual reactions
- Handled 65-95 time-steps of fluorescence readings per reaction

The high-throughput application makes QuICSeedR suitable for large-scale studies and high-volume data analysis in research settings.

```
#Read data with pre-defined helper function that format values as needed
grinder_data = BulkReadMARS(path = './data/grinder/',
                            plate_subfix = 'plate',
                            raw_subfix = 'raw',
                            helper_func = flip_and_replace)

#Set parameters in the workflow
params = list(
  CleanMeta = list(split_content = TRUE,
```

```
                   split_into = c("dilution", "sampleID")),
  GetCalculation = list(norm = TRUE, norm_ct = 'pos',
                        sd_fold = 10,
                        cycle_background = 6),
  GetAnalysis = list(control = 'neg', alternative = "greater"),
  SummarizeResult = list(sig_method = 'metric_count', method_threshold = 3)
)

#Get results
results = BulkProcessing(data = grinder_data, params = params)
```

In addition to adding metadata about the experimental setup, it is also easy to add existing metadata about study design after data analysis.

```
#Extract calculation of metrics and prepare the data for adding study metadata
result = results$combined_calculation
sel = c('blank', 'pos', 'neg')
sel = which(result$content %in% sel)
result = result[-sel, ]
colnames(result)[7] = 'SampleID'

#Add study metadata
metadata = read_xlsx('data/grinder_summary.xlsx')
full_result = right_join(metadata, result)
```

A subset of the 1,152 reactions is visualized here. Samples 10e1__2P through 10e3__5P represent swabs from equipment used to process CWD-negative muscle tissue. Samples 10e3__20P through 10e1__24P are from equipment that processed CWD-positive muscles, which demonstrated significant seeding activity in RT-QuIC assays. Lastly, samples 10e1__64P to 10e2__72P are swabs from equipment used to process CWD-positive brain samples.

```
col_palette <- viridis(50, option = 'turbo')

PlotMetric(data, x = 'replicate', y = 'content', fill_var = 'RAF',
           box = FALSE, point = FALSE) +
  geom_tile(color = "white",
            lwd = 1,
            linetype = 1) +
  scale_y_discrete(limits = rev(levels(factor(data$content))))+
  scale_fill_gradientn(colors = col_palette) +
  coord_flip() +
  theme(axis.text.x = element_text(angle = 45, hjust = 1),
        aspect.ratio = 1/5)
```

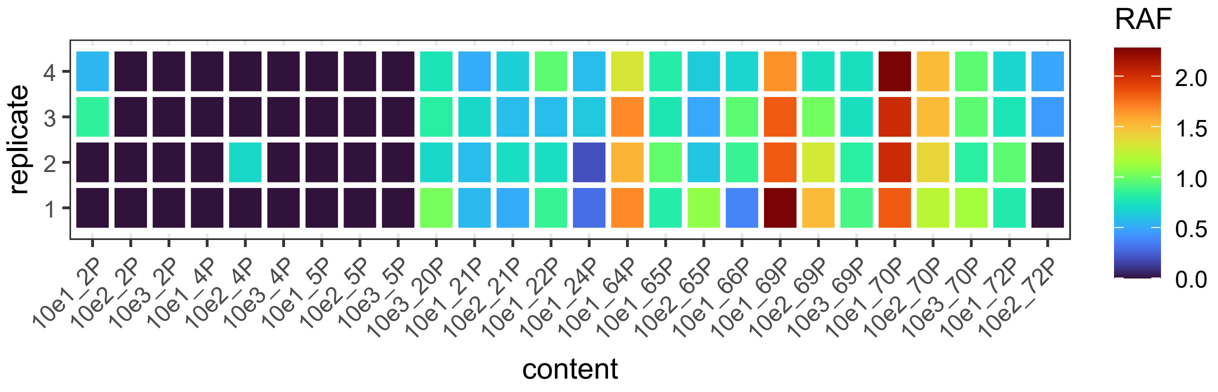


**2) Applications across various tissue types** In this section, QuICSeedR is applied to RT-QuIC data generated from various tissue types across different species, analyzing RT-QuIC data from blood and muscle samples from white-tailed deer (Schwabenlander et al., 2021; Li et al., 2021), as well as ear tissue samples from elk (Bryant et al., 2024). These diverse applications underscore QuICSeedR's flexibility in CWD research, benefiting data analysis for both CWD researchers and wildlife managers.

**a. Blood and muscle** In this demonstration, data from both blood and muscle samples are analyzed and visualized. Data was generated using samples obtained from the same group of ELISA-positive white-tailed deer, which were part of different studies (Schwabenlander et al., 2021; Li et al., 2021). For details of study and experiments, see orignal publications (Schwabenlander et al., 2021; Li et al., 2021).

```
#Load data
muscle_data = BulkReadMARS(path = './data/muscle/',
                           plate_subfix = 'plate', raw_subfix = 'raw')
blood_data = BulkReadMARS(path = './data/blood/',
                          plate_subfix = 'plate', raw_subfix = 'raw')

#Set parameters for analysis
params = list(
  GetCalculation = list(sd_fold = 5, time_skip = 5, cycle_background = 6),
  GetAnalysis = list(control = 'neg', alternative = "greater"),
  SummarizeResult = list(sig_method = 'RAF')
)
```

```r
#Get results
results_muscle = BulkProcessing(data = muscle_data, params = params)
results_blood = BulkProcessing(data = blood_data, params = params)

#Extract data calculating metric of interest (RAF) and remove name of the
#experiment
combined_calculation_muscle = results_muscle[['combined_calculation']][,-11]
combined_calculation_blood = results_blood[['combined_calculation']][,-11]

#ELISA positive animals
elisa_pos = c('166', '250', '333','353','360','363','376','384','508','735')

#Include 826, a muscle sample from an ELISA - animal
sel0 = c(elisa_pos, '826')
sel = which (combined_calculation_muscle$content %in%  sel0)
vis0 = combined_calculation_muscle[sel, ]
vis0$tissue = 'muscle'

#Include 777, a blood sample from an ELISA - animal
sel1 = c(elisa_pos,'777')
sel = which (combined_calculation_blood$content %in%  sel1)
vis1= combined_calculation_blood[sel,]
vis1$tissue = 'blood'

#Data visualization
visall = rbind (vis0, vis1)
visall$content = factor(visall$content, levels = c('777','826', sel0[1:10]))

PlotMetric(visall, y = 'RAF', fill_var = 'tissue', box = FALSE, point = FALSE,
   l1 = geom_boxplot(color = 'black', position = position_dodge(0.8)),
   l2 = geom_point(position = position_dodge(0.8)),
   l3 = scale_fill_brewer(palette="RdBu"),
   l4 = scale_y_continuous(limits = c(0,0.17)))
```

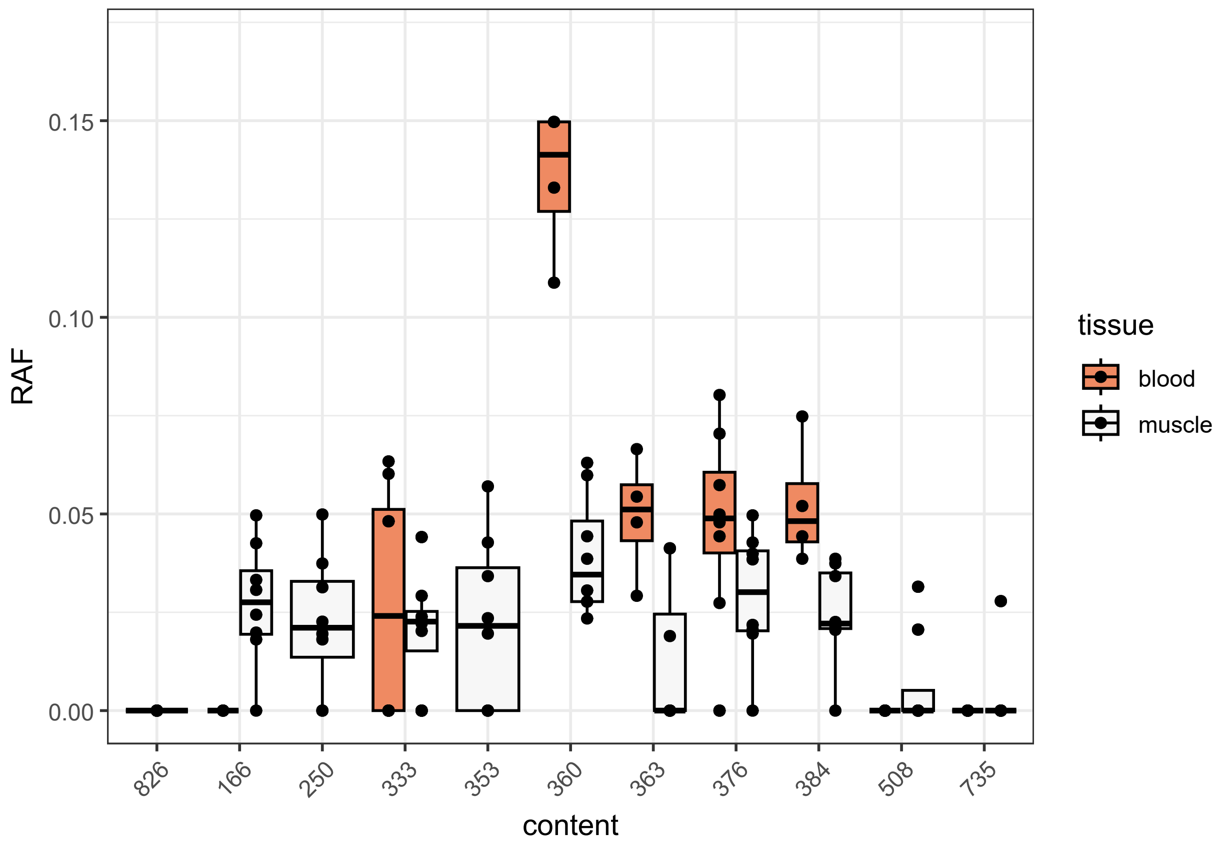


**b. Elk ear** For this demonstration, a subset of RT-QuIC data from elk ear tissue samples collected in the Black Hills and southwestern regions of South Dakota was utilized. These samples were acquired opportunistically from multiple sources, including cullings, hunters' harvests, predator kills, and naturally deceased elk found in the area (Bryant et al., 2024). From each intact ear, four distinct locations, designated as ventral (V), medial (M), dorsal (D), and lateral (L) were analyzed. For detailed information on the experimental procedures, please refer to the original publication by Bryant et al. (2024).

```
#Loading data, setting parameters for analysis, and getting results
add_underscore <- function(text) {
  gsub("([a-zA-Z])(\\d)", "\\1_\\2", text)
}

elkear = BulkReadMARS(path = './data/elkear', plate_subfix = 'plate',
                      raw_subfix = 'raw', helper_func = add_underscore)

params = list(
  CleanMeta = list(split_content = TRUE, split_into = c('region', 'sample')),
  GetCalculation = list(cycle_background = 5, norm = TRUE, norm_ct = 'Pos',
                        sd_fold = 10, time_skip = 5),
  GetAnalysis = list(control = "Neg")
)

results = BulkProcessing(data =elkear, params = params)

#Using percentage of replicates crossing the threshold as method to
```

```r
#determine whether a sample has seeding activity
kable(head(results[['combined_result']][,c(1:3, 12:14)]))
```

| content | result | method | xth_count | total_rep | xth_percent |
|---|---|---|---|---|---|
| bl | | xth_percent | 3 | 8 | 37.5 |
| Neg | | xth_percent | 0 | 8 | 0.0 |
| Pos | * | xth_percent | 8 | 8 | 100.0 |
| V_5 | * | xth_percent | 8 | 8 | 100.0 |
| M_5 | * | xth_percent | 8 | 8 | 100.0 |
| D_5 | * | xth_percent | 8 | 8 | 100.0 |

```r
#Data cleaning and visualization
calc = results[['combined_calculation']]
sel = c('bl', 'Pos', 'Neg')
sel = which(calc$content %in% sel)
calc = calc[-sel, ]

calc$sample = factor( calc$sample, levels = c('5', '15','16', '28', #ELISA +
                                              '6', '24', '29', '30')) #ELISA -

col_palette <- viridis(100, option = 'magma')

PlotMetric(calc, x = 'sample', y = 'region', fill_var = 'RAF',
           box = FALSE, point = FALSE)  +
  geom_tile(color = "white",lwd = 1, linetype = 1) +
  coord_fixed() +
  scale_fill_gradientn(colors = col_palette)
```

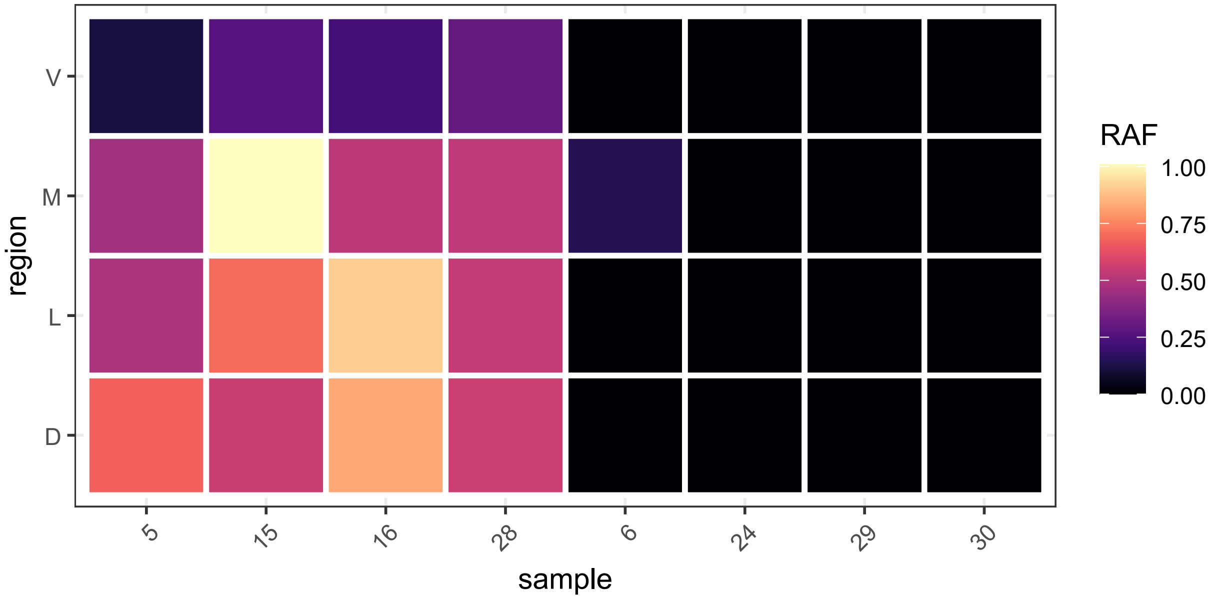
V
M
L
D
region
5
15
16
28
6
24
29
30
sample
RAF
1.00
0.75
0.50
0.25
0.00

**3) Human neurodegenerative diseases applications** Finally, QuICSeedR was applied to human neurodegenerative diseases by analyzing existing nano-QuIC and F-PMCA data related to synucleinopathies. The detection of misfolded alpha-synuclein is of significant clinical value in these disorders. This application demonstrated QuICSeedR's effectiveness in enhancing the analysis of F-SAA data for protein misfolding in human disease contexts.

**a. Detecting misfolded alpha-synuclein seeding activity in human plasma from Parkinson's disease (PD) patients using nano-QuIC** Nano-QuIC significantly accelerates and improves the sensitivity of misfolded alpha-synuclein detection (Christenson et al., 2023). The dataset used here comprises Nano-QuIC results from individual patient plasma samples generously provided by the Mayo Clinic. Refer to the original publication for details of patient information and experiments (Christenson et al., 2023).

Using a cut-off time of approximately 250 hours as well as criteria consisting of a fluorescence threshold of 5000 relative fluorescence units (RFUs) and 50% replicates crossing the threshold, 3 out of 4 patients with idiopathic PD could be identified.

```
#Read in data
plasma_data = BulkReadMARS(path = './data/plasma',
                           plate_subfix = 'plate',
                           raw_subfix = 'raw')

#Set up parameters for analysis
params = list(
  CleanMeta = list(split_content = TRUE,
                   split_into = c("condition", "ID")),

  #Set cycles for cut-off
  CleanRaw = list(cycle_total = 325),

  #Use 5000 relative fluorescence unit as the threshold
  GetCalculation = list(threshold_method = 'rfu_val', rfu = 5000)
)

#Get results
results = BulkProcessing(data = plasma_data, do_analysis = FALSE,
                         params = params)

#Summarized results
kable(results[['combined_result']][,1:7])
```

| content | result | method | position | xth_count | total_rep | xth_percent |
|---|---|---|---|---|---|---|
| HC_1 | | xth_percent | D01-E01-F01-G01-H01 | 0 | 5 | 0 |
| HC_2 | | xth_percent | D02-E02-F02-G02-H02 | 0 | 5 | 0 |
| PD_15 | * | xth_percent | D03-E03-F03-G03-H03 | 4 | 5 | 80 |
| PD_16 | * | xth_percent | D04-E04-F04-G04-H04 | 3 | 5 | 60 |
| HC_3 | | xth_percent | A01-B01-C01-D01 | 0 | 4 | 0 |
| HC_4 | | xth_percent | A02-B02-C02-D02 | 0 | 4 | 0 |
| PD_17 | | xth_percent | A03-B03-C03-D03 | 0 | 4 | 0 |
| PD_22 | * | xth_percent | A04-B04-C04-D04 | 3 | 4 | 75 |

Visualization in the original paper is also recreated using QuICSeedR (Christenson et al., 2023).

```r
#Data visualization
col1 = c('black', 'gray', 'pink','red')
col2 = c('darkblue', 'lightblue', 'orange','brown')

par(mfrow = c(1,2))
PlotRawMulti(results$combined_cleanraw$`20240415_r12`,
             samples = c('HC_1', 'HC_2', 'PD_15', 'PD_16'),
             ylim = c(0,85000),
             custom_colors = col1)

PlotRawMulti(results$combined_cleanraw$`20240528_r12`,
             samples = c('HC_3', 'HC_4', 'PD_17', 'PD_22'),
             ylim = c(0,85000),
             custom_colors = col2)
```

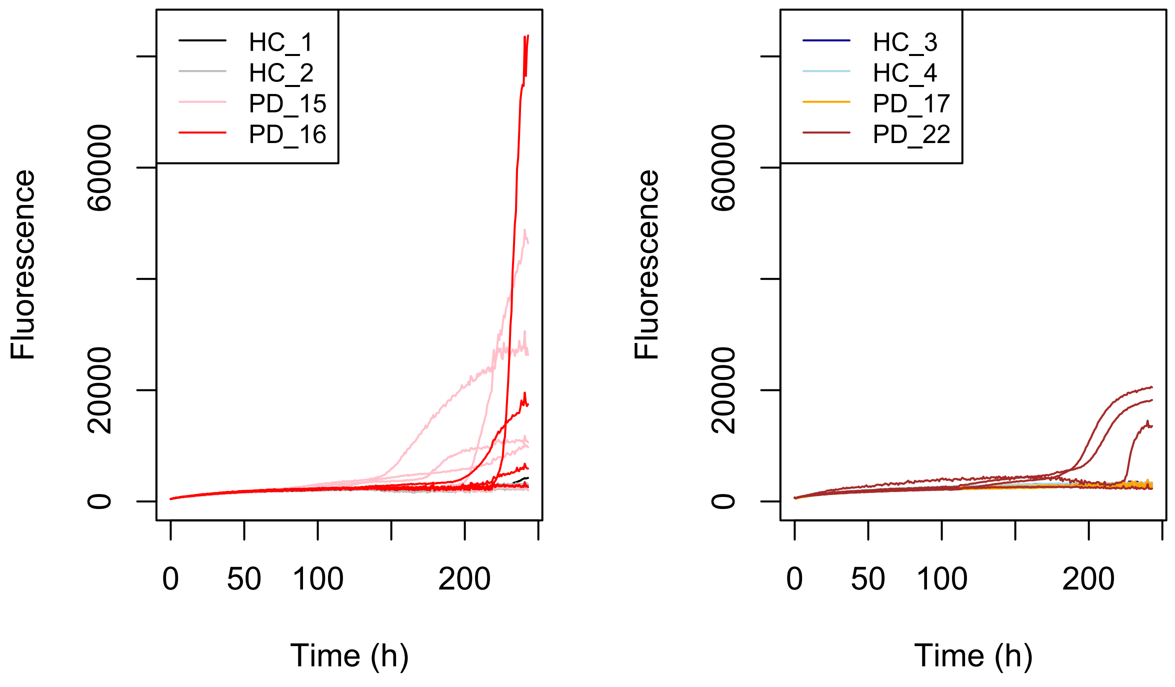


```r
par(mfrow = c(1, 1))
```

**b. Detecting misfolded alpha-synuclein seeding activity in human cerebrospinal fluid (CSF) from PD and multiple system atrophy (MSA) patients using F-PMCA.** F-PMCA is widely adopted for detecting alpha-synuclein seeding activity in synucleinopathies. Shahnawaz et al. (2020) utilized fluorescence intensity in F-PMCA to differentiate CSF samples from individuals with Parkinson's disease (PD), multiple system atrophy (MSA), and controls. To demonstrate the application of QuICSeedR to F-PMCA data, QuICSeedR was applied to fluorescent data from Shahnawaz et al. (2020), which includes CSF samples from 42 control individuals, 30 MSA patients, and 47 PD patients. For detailed information on patient selection criteria, experimental design, and data generation methods, please refer to the original study by Shahnawaz et al. (2020).

```
#Load data
data_Shahnawaz2020 = read_xlsx('./data/Shahnawaz_2020.xlsx')
time = data_Shahnawaz2020[["Time (h)"]]

#format data to use with QuICSeedR analysis
raw_Shahnawaz = as.matrix(data_Shahnawaz2020[,-1])
rownames(raw_Shahnawaz) = time

meta_Shahnawaz = data.frame(content = colnames(raw_Shahnawaz),
                            condition = gsub("[^A-Za-z]", "", colnames(raw_Shahnawaz)))

#Plotting a subset of samples
cols = c(1:10, 43:52, 73:82)
PlotRawMulti(raw = raw_Shahnawaz[,cols], c('HC','MSA','PD'),
             custom_colors =  c("gray", "pink", '#c1121f'))
```

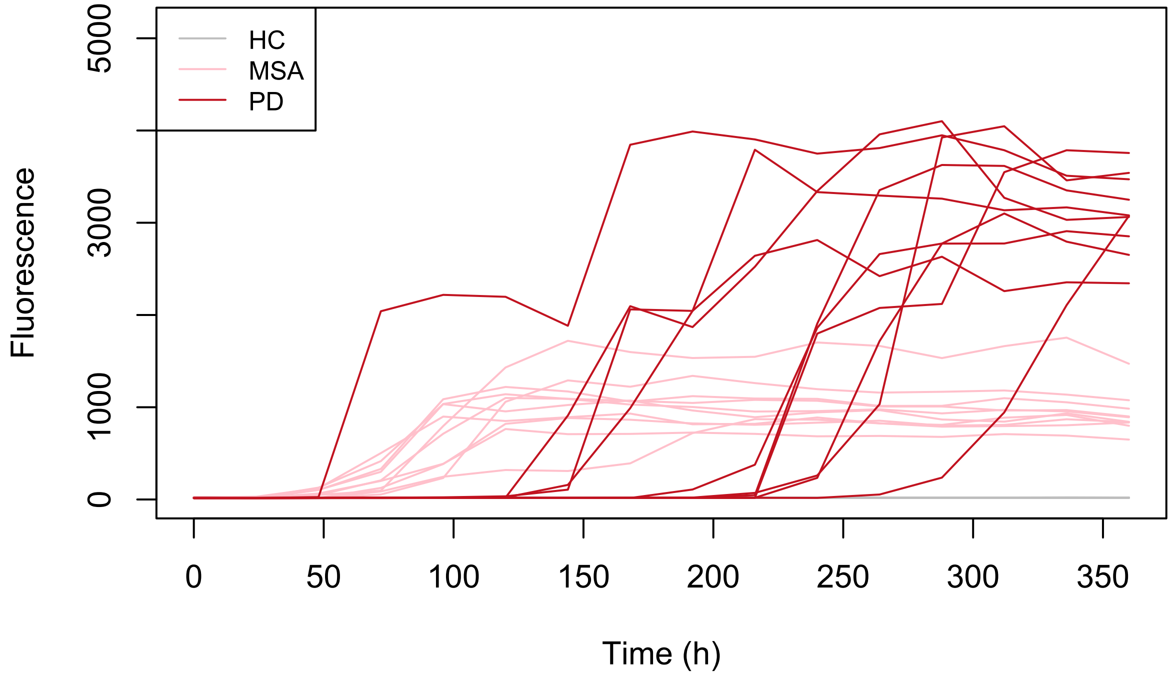

Minimizing the degree of overlap is critical for achieving the specificity necessary to accurately distinguish between PD and MSA. Utilizing existing metrics, including those integrated within the package, and the area under the curve (AUC) used by Russo et al. (2021), the performance of each metric was evaluated. This evaluation exemplified QuICSeedR's capability in generating reproducible and comparative analyses of these metrics.

```r
#Calculation by QuICSeedR
calc_Shahnawaz = GetCalculation(raw_Shahnawaz, meta_Shahnawaz,
                                time_skip = 1, cycle_background = 2, binw = 10)

#Calculate AUC
library(pracma)
calc_Shahnawaz$AUC = apply(raw_Shahnawaz, 2, function(col) {
  trapz(time, col)
})
```

Wilcoxon rank-sum tests were then conducted for each metric and extracted the p-values, which all indicated significant differences among conditions. However, the graphs revealed varying levels of separation between conditions.

```r
kable(p_values_table)
```

| Metric | HC_vs_MSA | HC_vs_PD | MSA_vs_PD |
|---|---|---|---|
| RAF | 3.52e-16 | 9.07e-18 | 1.69e-11 |
| MS | 1.22e-20 | 4.40e-26 | 9.45e-22 |
| MPR | 1.22e-20 | 4.40e-26 | 3.35e-17 |
| AUC | 1.22e-20 | 4.40e-26 | 4.36e-12 |

```r
#Wrapper function for PlotMetric() to reduce code redundancy
create_plot <- function(data, y_var) {
  PlotMetric(data, x = 'condition', y = y_var,
             fill_var = 'condition', box = FALSE, point = FALSE) +
    geom_boxplot(aes(x = factor(calc_Shahnawaz$condition)),
                 outlier.shape = NA) +
    geom_jitter(width = 0.2, height = 0, size = 1) +
    xlab('Condition') +
    scale_fill_manual(values = c("gray", "pink", '#c1121f'))
}

P1 <- create_plot(calc_Shahnawaz, 'RAF')
P2 <- create_plot(calc_Shahnawaz, 'MS')
P3 <- create_plot(calc_Shahnawaz, 'MPR')
P4 <- create_plot(calc_Shahnawaz, 'AUC')

#Visualizing separation of conditions by each metric
grid.arrange(P1, P2, P3, P4, nrow = 2, ncol = 2)
```

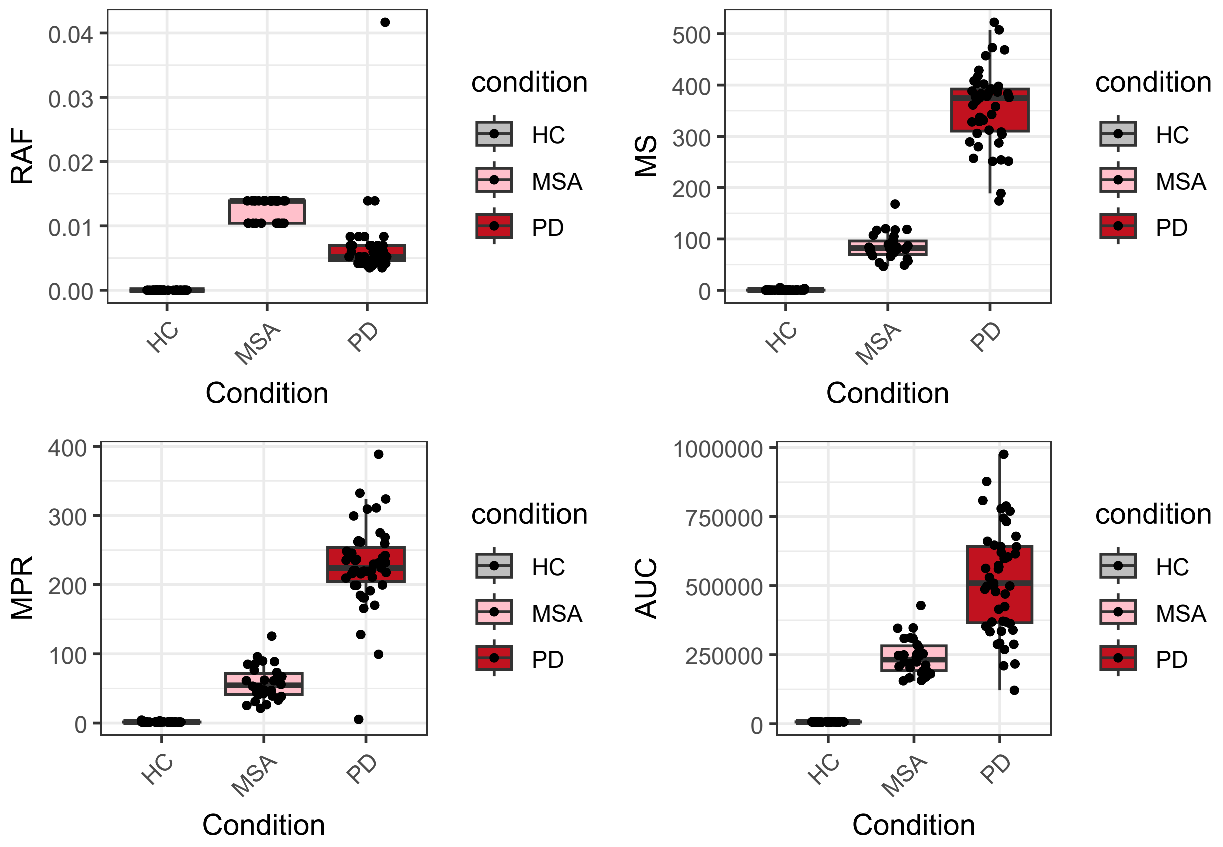


To quantify the differences observed in the ability of each metric to separate conditions, the degree of overlap between conditions for each metric was measured. This involved determining how many PD values for each metric fall within the range of MSA values, how many MSA values fall within the range of PD values, and how many control values fall within the ranges of both PD and MSA values, and vice versa. By analyzing these overlaps, the metrics' effectiveness in distinguishing conditions, especially between PD and MSA, can be better characterized.

Based on the calculation above, MS (maximal rate of change of fluorescence over 240 hours) can 100% differentiate PD and MSA. Therefore, the MS metric, along with further research into metric performance, may enhance future diagnostic value by achieving higher specificity and sensitivity.

```r
metrics <- c('RAF', 'MS', 'MPR', 'AUC')
results <- do.call(rbind, lapply(metrics, function(metric)
  count_values_in_range(calc_Shahnawaz, metric)))

kable(results)
```

| Metric | HC_in_MSA | HC_in_PD | MSA_in_HC | PD_in_HC | PD_in_MSA | MSA_in_PD |
|---|---|---|---|---|---|---|
| RAF | 0 | 0 | 0 | 0 | 2 | 30 |
| MS | 0 | 0 | 0 | 0 | 0 | 0 |
| MPR | 0 | 0 | 0 | 0 | 1 | 30 |
| AUC | 0 | 0 | 0 | 0 | 16 | 30 |
