## Supplementary figures and images for "QuICSeedR: An R package for analyzing fluorophore-assisted seed amplification assay data"

### grinder_MPR.png

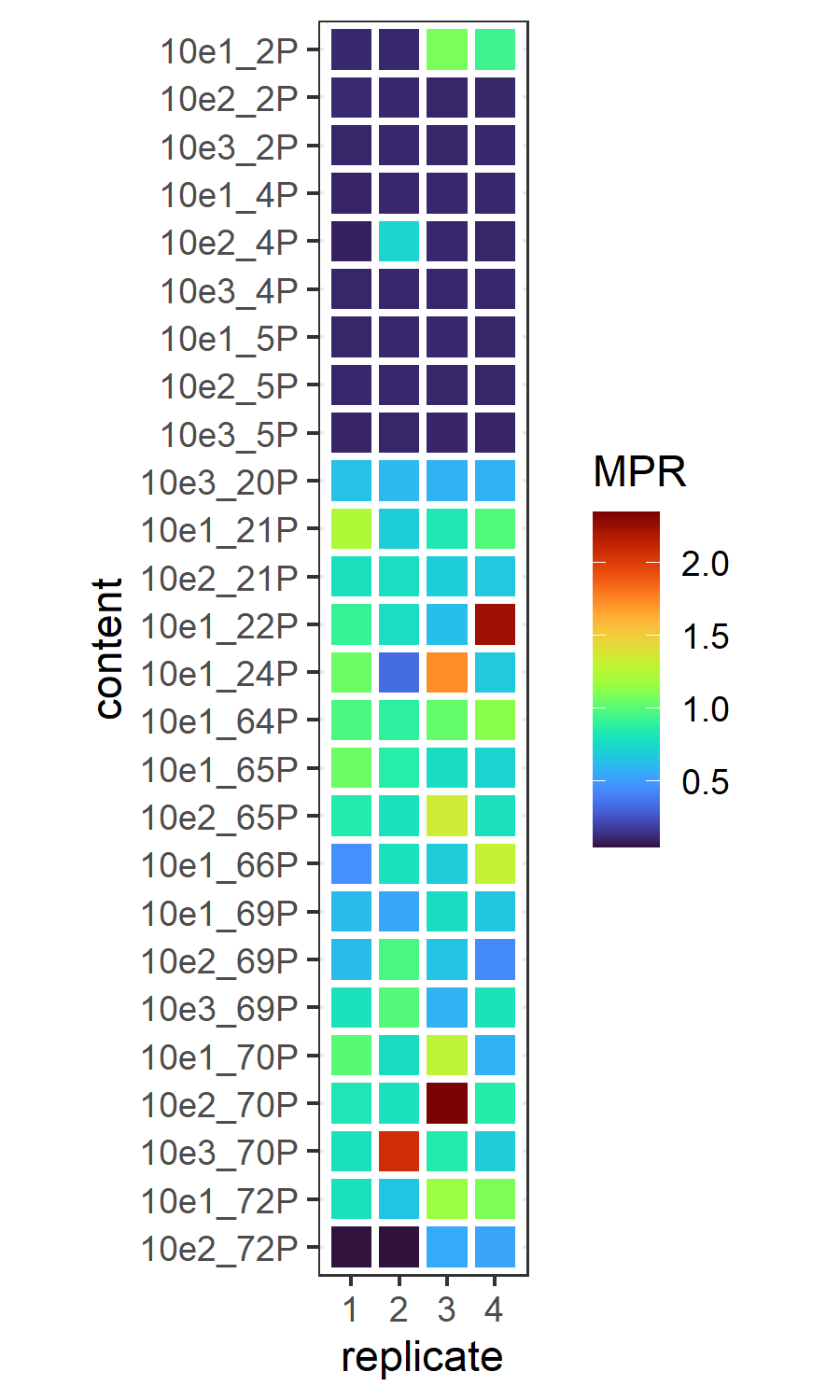

### grinder_MS.png

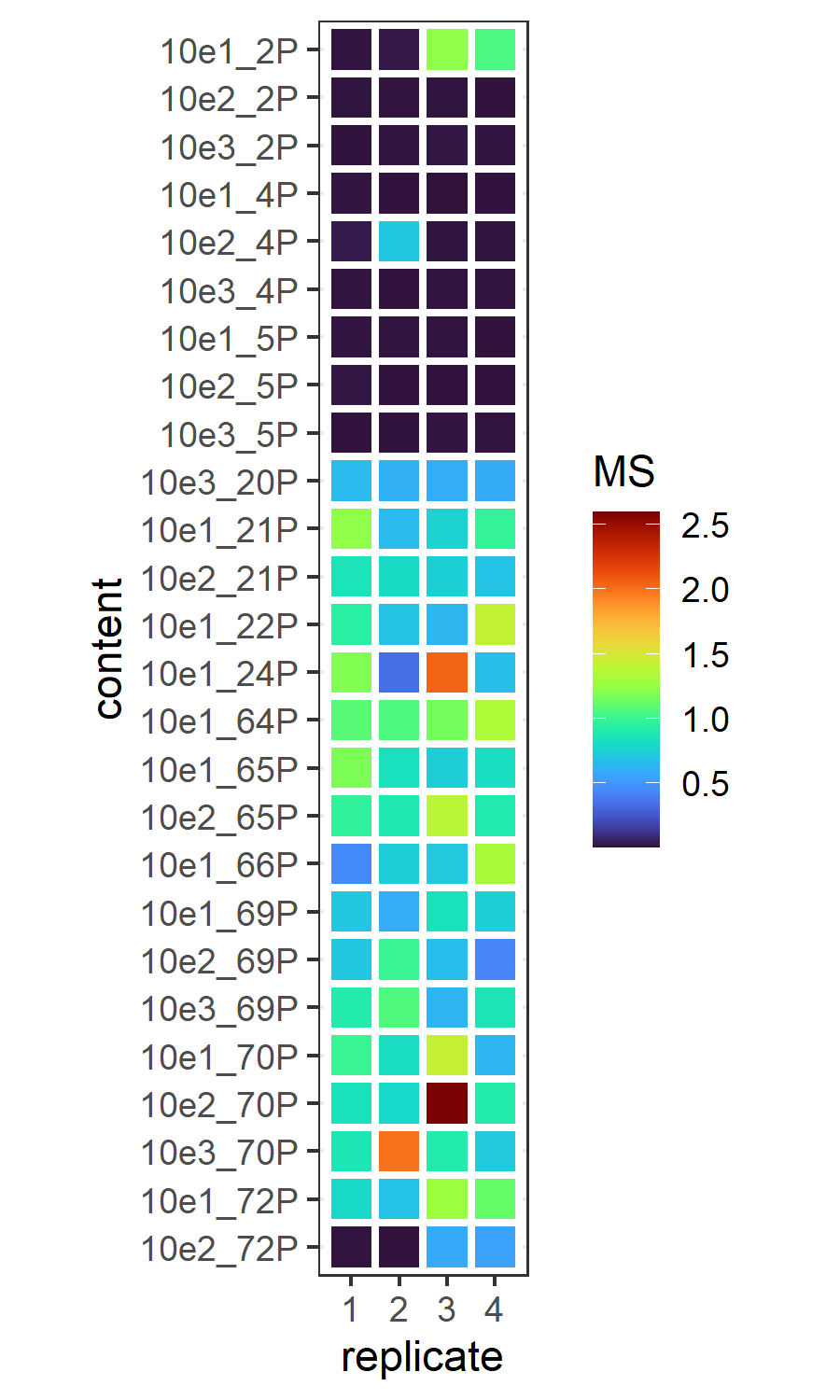

### grinder_RAF.png

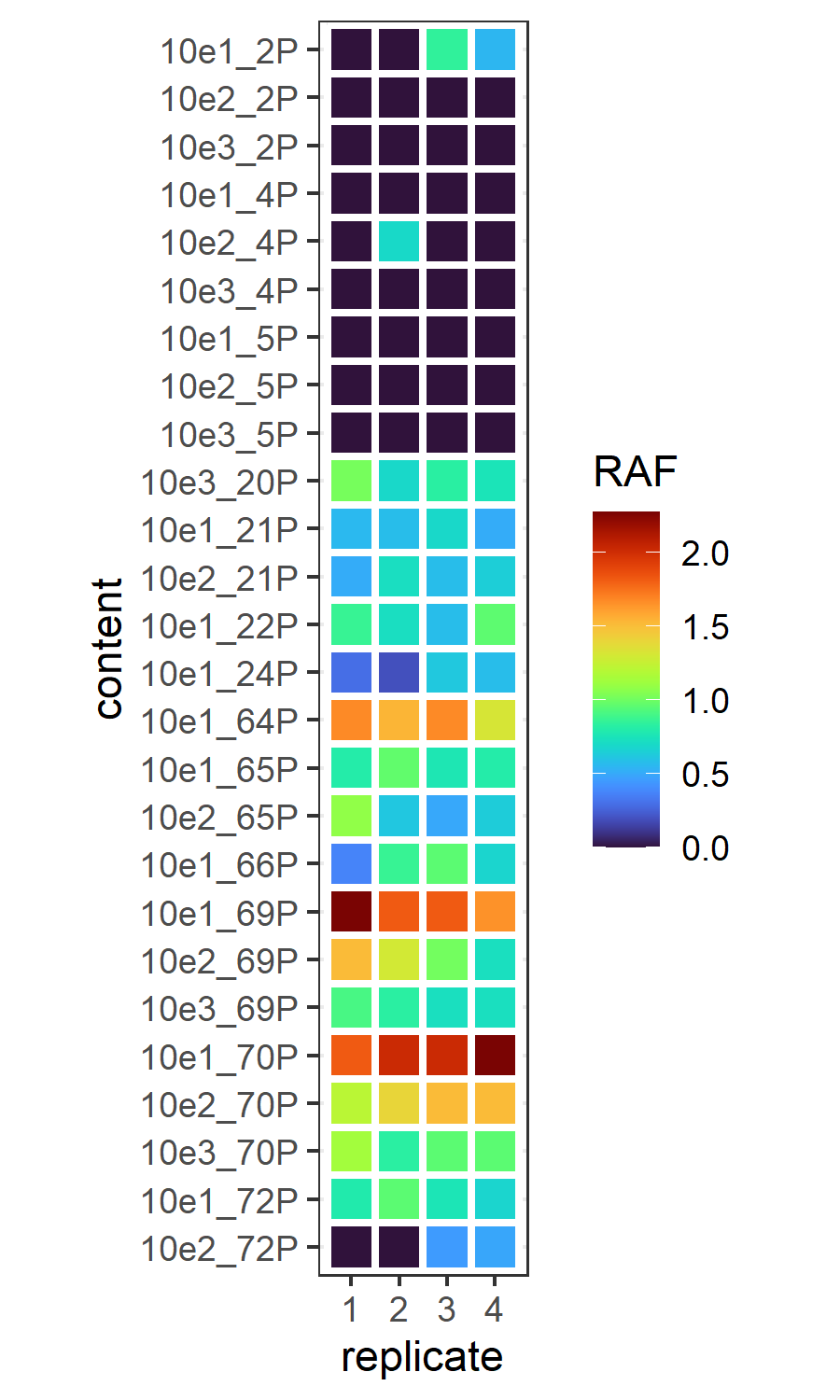

### Screenshot 2024-07-16 120241.png

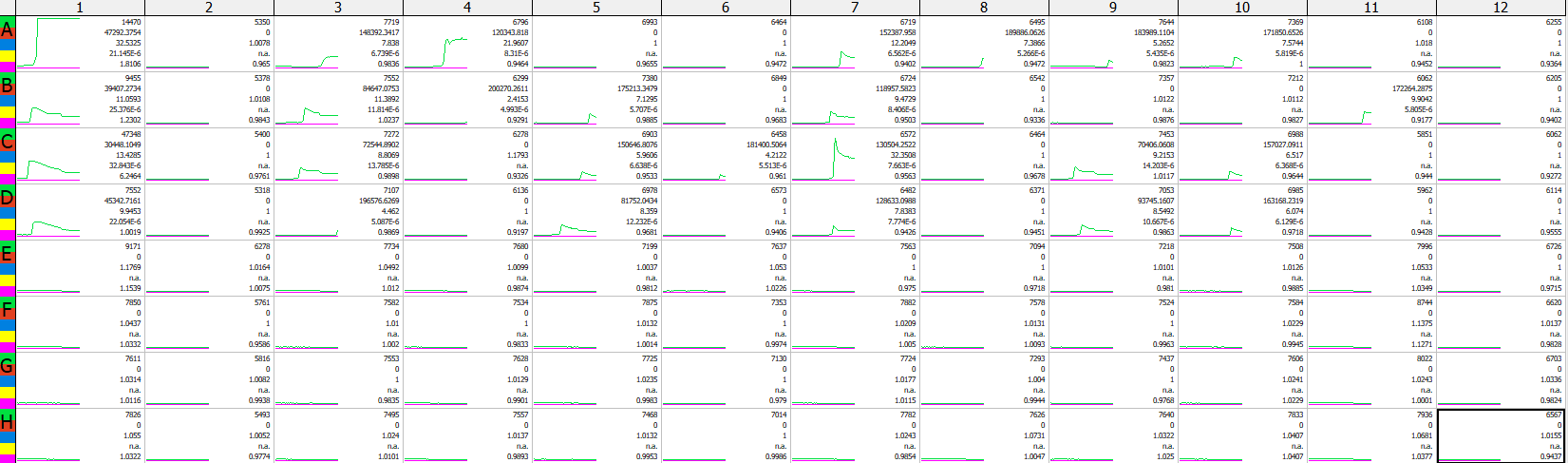

### Screenshot 2024-07-16 120409.png

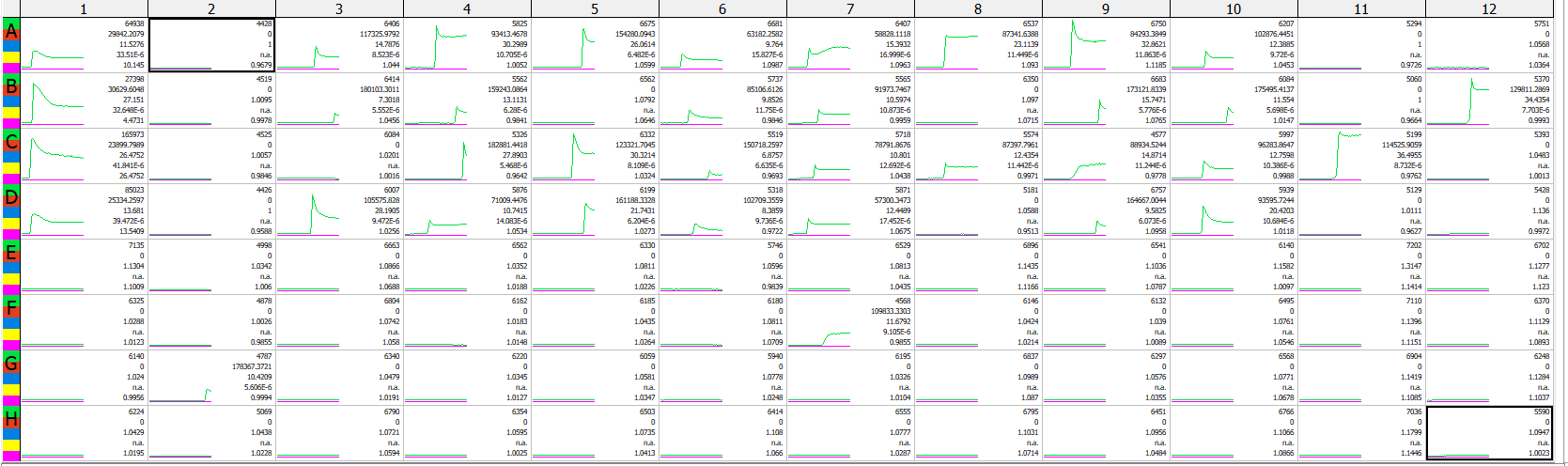
